# Cachexia and fibrosis are costs of chronic IL-1R-mediated disease tolerance in *T. gondii* infection

**DOI:** 10.1101/783316

**Authors:** Stephanie J. Melchor, Jessica A. Hatter, Erika A. LaTorre Castillo, Claire M. Saunders, Kari A. Byrnes, Imani Sanders, Daniel Abebayehu, Thomas Barker, Sheryl Coutermarsh-Ott, Sarah E. Ewald

## Abstract

Cachexia is an immune-metabolic disease of progressive muscle wasting that impairs patient survival and quality of life across a range of chronic diseases. *T. gondii* is a protozoan parasite that causes lifelong infection in many warm-blooded organisms, including humans and mice. Here we show that mice infected with *T. gondii* develop robust, sustained cachexia and perivascular fibrosis in metabolic tissues. Consistent with an emerging role for the IL-1 axis in disease tolerance, we show that mice deficient in the Type 1 IL-1 receptor (IL-1R) have more severe acute muscle wasting, adipocyte and hepatocyte necrosis, independent of parasite burden. Unexpectedly, IL-1R^-/-^ mice rapidly recover from acute disease, despite sustained parasite infection, and are protected from chronic cachexia as well as perivascular liver and muscle fibrosis. These data are consistent with a model where IL-1R signaling benefits cell survival and tissue integrity over short periods of inflammation, but sustained reliance on IL-1 mediated tolerance programs come at the cost of fibrosis and cachexia.

**Summary:** IL-1R signaling drives a disease tolerance program that protects mice from tissue pathology during acute *Toxoplasma gondii* infection. However, extended IL-1R signaling drives chronic cachexia and perivascular fibrosis in the liver and skeletal muscle.

## Introduction

Cachexia is defined as the loss of 5% of lean body mass in under six months accompanied by at least three of the following symptoms: weakness, fatigue, adipose loss, abnormal blood biochemistry and/or anorexia (Evans *et al*., 2008). In industrialized countries (United States, Europe and Japan) where the frequency of cachexia is well-documented, approximately 1% of the population or 9 million patients are cachectic (von Haehling, Anker and Anker, 2016). As a disease that occurs in conjunction with a primary chronic disease, the prevalence of cachexia can range from 5-15% in chronic heart failure and chronic pulmonary disease to over 80% in advanced cancer patients (Suzuki *et al*., 2013; von Haehling and Anker, 2014). Cachexia impairs patient quality of life, limits the effectiveness and duration of therapeutic interventions and can directly cause death in chronically ill patients. Currently, there are no broadly efficacious therapeutics for cachexia. Dietary supplementation, anabolic steroids and TNF blocking treatments have largely failed in the clinic (Tisdale, 1997).

Our poor understanding of the molecular and physiological mechanisms of cachexia can be linked to limited animal models that accurately recapitulate the long-term progression of disease. Surgical interventions including cardiac, gastric and renal obstructive surgeries are lethal after a rapid period of weight loss; tumor models take longer to develop but have a similar 1-2 week window of weight loss before tumor growth is lethal; low level endotoxin injection is transient and in a new LCMV model of cachexia mice recover weight once infection is cleared (Deboer, 2009; Baazim *et al*., 2019). Cachexia or cachexia-like symptoms have been studied in a range of parasite infection settings; however, these have received little attention in the modern molecular study of disease (Tacey and Cerami, 1989).

We have recently shown that per oral infection with the protozoan parasite *Toxoplasma gondii* (*T. gondii*) leads to a sustained loss of muscle mass that meets the current clinical definition of cachexia (Hatter *et al*., 2018). *T. gondii* is an obligate intracellular parasite that infects a wide range of mammalian hosts in nature, including mice and humans. Infection occurs when a host ingests food contaminated with parasite tissue cysts or oocysts shed from a feline definitive host (Montoya and Liesenfeld, 2004). *T. gondii* invades the small intestine and disseminates throughout the host as the rapidly replicating tachyzoite form. Over the first two weeks of infection the parasite can be detected in almost every tissue in the body (Di Cristina *et al*., 2008). A TH1-mediated immune response is necessary to restrict infection and eventually clears systemic parasitemia; however, the host remains infected for life with low titers of *T. gondii* predominantly in the bradyzoite cyst form in the brain, muscle and other tissues (Robert-Gangneux F *et al*., 2018). An active immune response is critical for chronic suppression of the parasite as evident in sustained, high titers of serum innate inflammatory cytokines, *T. gondii*-specific IgG and the observation that parasites recrudesce in immune suppressed patients (Pernas *et al*., 2014; Robert-Gangneux F *et al*., 2018). The innate immune cytokines TNF-α, IL-1α, IL-6 and IFN-γ also comprise an inflammatory signature frequently observed in cachectic patients,(de Matos-Neto *et al*., 2015) and administration of recombinant TNF-α, IL-1α, IL-6 and IFN-γ is sufficient to induce muscle atrophy and/or cachexia in rodent models (Oliff *et al*., 1967; Fong *et al*., 1989; Matthys *et al*., 1991; Plata-Salaman *et al*., 1997; Haddad *et al*., 2005). Despite the strong historical association of TNF-α (previously called cachectin) with cachexia, efforts to block it in the clinic have not been effective using etanercept, which blocks TNF-α signaling or infliximab, a TNF-α blocking antibody (Goldberg *et al*., 1995; Marcora, Chester and Mittal, 2006; Jatoi *et al*., 2010). In contrast, a first-in-class human monoclonal antibody against IL-1α was recently shown to reduce pain and fatigue while improving weight gain in metastatic cancer patients suggesting that targeting IL-1 signaling during cachexia may be more effective (Hong *et al*., 2014). A role for IL-1 has also recently emerged as a regulator of disease tolerance programs in endotoxic shock and in *Salmonella typhimurium* infection (Rao *et al*., 2017; Benjamin *et al*., 2018). Disease tolerance programs protect the host from bystander damage during infection without directly affecting pathogen load, whereas resistance programs improve host fitness by targeting pathogenic organisms for clearance (Medzhitov, Schneider and Soares, 2012). Taken together, these findings suggest that the IL-1 axis regulates fundamentally distinct biology than canonical pro-inflammatory signaling cascades.

Given the robust and highly reproducible nature of *T. gondii* infection-induced cachexia, we asked if this model could provide novel insight into the pathophysiology of cachexia. Although cachexia in our per oral model was stable even after intestinal inflammation had resolved, we wanted to know if bypassing the intestinal tract entirely could still recapitulate the cachectic diseased state. Here we report that bypassing the small intestine by infecting mice via the intraperitoneal route was sufficient to induce chronic cachexia. IL-1 signaling was necessary to protect mice from tissue necrosis during acute cachexia; however, immune infiltrate and parasite load was similar between wildtype and IL-1R knockout mice consistent with a role for IL-1 in tolerance to inflammation rather than directing a parasite restriction mechanism. Unexpectedly, after the first four weeks of infection, IL-1R deficient mice rapidly recovered from acute weight loss, including muscle atrophy. This was not due to differences in chronic parasite load. The longevity of our model revealed that cachectic mice develop perivascular fibrosis in major metabolic tissues including the visceral white adipose tissue, the skeletal muscle and the liver which was also dependent on IL-1R signaling. Cumulatively, these data suggest that while IL-1 signaling can be beneficial during the acute inflammatory response to *T. gondii*, sustained reliance on IL-1-mediated tolerance biology can have maladaptive consequences for host fitness.

## Results

### *T. gondii* infection leads to sustained cachexia in mice

We previously demonstrated that mice infected with *T. gondii* via the per oral route develop cachexia, which was sustained even after intestinal inflammation resolved. This prompted us to ask whether *T. gondii* infection could induce cachexia if the gastrointestinal tract was bypassed using intraperitoneal infection (Evans *et al*., 2008; Hatter *et al*., 2018). 10-14 week old male C57BL/6J mice were intraperitoneally injected with 10 bradyzoite tissue cysts of the Type II *T. gondii* strain Me49. *T. gondii*-infected mice lost 15-20% of their initial body mass during the first three weeks of infection (Fig 1A-B), when parasites are systemic. After the first three weeks of infection, systemic parasitemia (parasites in the blood) resolves and *T. gondii* is driven into chronic tissue cyst form (Saeij *et al*., 2005; Djurković-Djaković *et al*., 2012). However, mice were unable to regain weight to the level of uninfected controls for up to twenty-one weeks (Fig. 1B), the duration of the experiment. Chronic infection was confirmed by counting cysts in brain homogenate using morphology and dolichos biflourus agglutinin staining of the cyst wall at nine weeks post infection (Fig. 1C). Infected mice went through a period of acute anorexia 8-10 days post-infection; however, they regained eating relative to uninfected controls indicating that sustained weight loss was not due to prolonged anorexia. In fact, infected mice trended towards eating more chow per 24-hour period than uninfected mice, although this was not significant (Fig. 1D, Supp. Fig 1A). Bomb calorimetry on fecal pellets confirmed that infected mice were absorbing similar number of calories as uninfected mice (Supp. Fig 1A).

**Figure 1:**
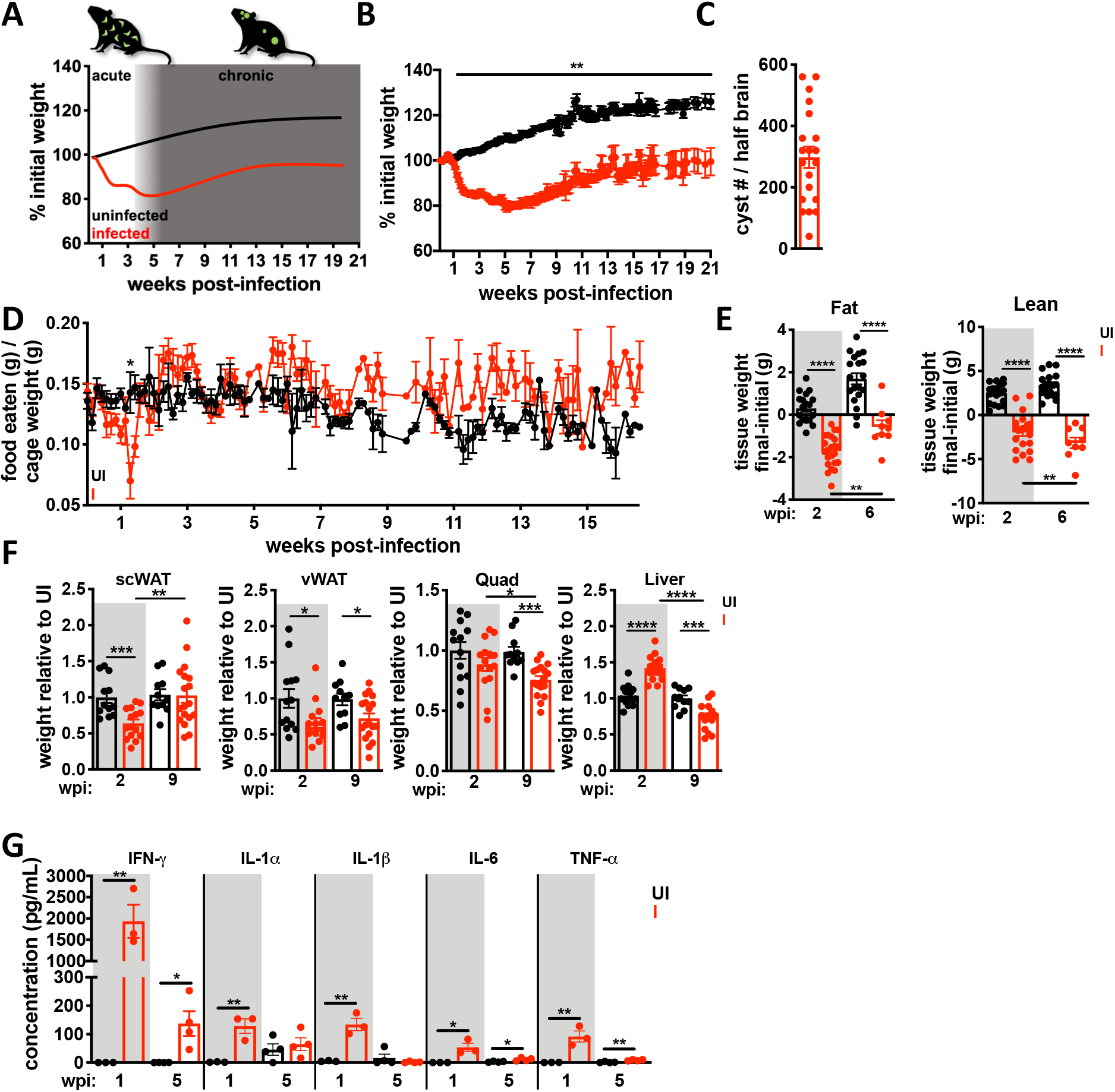
Chronic *Toxoplasma* infection causes sustained cachexia in mice. 10-14 week old C57BL/6J mice were intraperitoneally infected with 10 *Toxoplasma* cysts of the Me49-GFP-luciferase strain (I, red) or mock injected with PBS (UI, black). **A**, Schematic of weight loss relative to parasite distribution. The acute phase of infection (white) is dominated by *Toxoplasma* tachyzoites (green crescents) which spread systemically infecting most tissues in the body. 4-6 weeks post-infection, systemic infection is largely cleared and parasites are driven to the chronic tissue cyst form (green circles). **B**, Mice lose up to 20% of their initial body mass in the first 4 weeks of infection (I, red) and fail to regain weight relative to uninfected controls (UI, black). N=35-45 mice pooled from 3 independent experiments. **C**, Dolichos biflorous positive *Toxoplasma* cysts per half brain at 5-9 weeks post-infection. N=19 pooled from 6 independent experiments. **D**, Daily food intake per cage normalized to pooled weight of the mice in the cage measured every 24 hours. N=7-9 cages per group, pooled from 3 independent experiments. **E**, Echo MRI quantification of fat (left) and lean (right) tissue mass at 2 or 6 wpi. N=28-45 mice per group, representative of 3 independent experiments. **F**, Inguinal subcutaneous white adipose tissue (scWAT), epididymal visceral white adipose tissue (vWAT), quadriceps (Quad) and liver weights relative to the mean weight of uninfected tissue at 2 wpi or 9wpi N=12-18 mice per group, pooled from 2 experiments. **G**, Serum cytokines measured by Luminex at 1 or 5 wpi. N=3-4 mice per group, representative of 2 experiments. Error bars are standard error of the mean. *P < 0.05; **P < 0.01; ***P < 0.001, ****P < 0.0001 by unpaired Student’s T test with Holm-Sidak method to correct for multiple comparisons.

Loss of lean muscle mass is the primary diagnostic marker of cachexia. By two weeks postinfection, infected mice had a significant reduction in both fat (Fig. 1E, left) and lean (Fig. 1E, right) body mass by EchoMRI Whole Body Composition Analysis. Importantly, lean body mass wasting progressed by six weeks post-infection whereas fat body mass partially recovered (Fig. 1E). When individual tissues were dissected and weighed, we found that inguinal subcutaneous white adipose tissue (scWAT), epigonadal visceral white adipose tissue (vWAT) and quadricep muscle mass (Quad) were significantly reduced at two-weeks post infection and hepatomegaly was also observed, consistent with published reports of acute *T. gondii* infection in the liver (Atmaca *et al*., 2013). (Fig. 1F). Gastrocnemius, tibialis anterior and extensor digitorus longum muscles were also weighed and similarly reduced in mass (data not shown). By nine-weeks post-infection scWAT had recovered but vWAT, quad and liver were significantly reduced in size relative to uninfected littermate controls (Fig. 1F).

Consistent with fatigue associated with cachexia, mice had a significant reduction in night time activity four to five weeks post-infection measured by individually housing animals in CLAMS metabolic monitoring cages (Supp. Fig. 1B-F). CLAMS cages are commonly used to determine whether an animal relies on beta-oxidation or glycolysis as a primary energy source by measuring the rate of carbon dioxide production relative to oxygen consumption known as the respiratory exchange ratio (RER). We observed a significant depression in RER (Supp. Fig. 2B). This likely reflects a reduced demand for oxygen based on inactivity of infected mice rather than reliance on beta-oxidation as an energy source (Kir *et al*., 2014; Petruzzelli *et al*., 2014; Kliewer *et al*., 2015). Consistent with this conclusion, no significant differences in carbon dioxide production (VCO_2_, Supp. Fig. 1E) or oxygen consumption (VO_2_, Supp. Fig. 1D) were observed. This conclusion was also supported by significant reduction in the calculated production of heat by these mice (Supp. Fig. 1F). Although body temperature was significantly reduced at peak acute infection in infected mice, there was not a sustained difference during chronic infection between infected and uninfected mice as measured by subcutaneous temperature probes (Supp. Fig. 2A).

Elevated circulating inflammatory cytokines are a hallmark of cachexia that contribute to disease pathology (Morley, Thomas and Wilson, 2006). At one week post-infection, mice had elevated circulating IFN-γ, IL-1α, IL-1β, IL-6, and TNF-α compared to uninfected controls (Fig. 1G, grey background). Although TNF-α, IL-6, and IFN-γ were reduced by five weeks post-infection, they were still significantly elevated relative to uninfected controls (Fig. 1H, white background). IL-1α and IL-1β were not elevated in serum 5 weeks post-infection. This was expected because sustained systemic levels of these so-called pyrogens are a signature of shock, and it is well established that local levels of IL-1 can remain elevated without entering circulation (Dinarello, 2009; Bertheloot and Latz, 2017). Cumulatively, intraperitoneal *T. gondii* infection causes the progressive loss of muscle, loss of adipose tissue, transient anorexia, fatigue and elevated innate circulating cytokines consistent with the current, clinical definition of cachexia (Evans *et al*., 2008).

### *T. gondii*-induced cachexia causes perivascular fibrosis in metabolic tissues

Having established a robust and sustained form of cachexia during i.p. *T. gondii* infection, we next asked if we could discern long-term effects of cachexia on tissue homeostasis. We first asked if there was evidence of non-shivering thermogenesis or fat browning in our mice based on several reports using acute cancer cachexia models (Tsoli *et al*., 2012; Kir *et al*., 2014; Petruzzelli *et al*., 2014). Body temperature (Supp. Fig. 2A) and brown adipose depot size (Supp. Fig. 2B) were not significantly different between infected and uninfected mice at five weeks post-infection. There was a trend towards upregulation of *ucp1* (which uncouples mitochondrial respiration from ATP synthesis and drives thermogenesis) in scWAT and vWAT; however, *prdm16* a regulator of brown fat differentiation was downregulated in scWAT and other transcripts associated with browning were not significantly altered (Supp. Fig. 2C-E). The exception was *pparg*, which was significantly downregulated in scWAT and vWAT (Supp. Fig 2C-D). From these data we conclude that changes in brown fat distribution may occur in acute cachexia; however, non-shivering thermogenesis is not a main driver of chronic *T. gondii*-induced cachexia (Hatter *et al*., 2018).

Blocking lipolysis has been shown to limit muscle loss early in the onset of cachexia in cancer models (Arner and Langin, 2014; Kliewer *et al*., 2015). We assessed lipolysis machinery in the white adipose depots, muscle and liver at nine weeks post-infection by western blot. Although there was some variability between mice, we observed no consistent differences in the level of hormone-sensitive lipase (HSL), phosphorylated HSL, AKT, adipose triglyceride lipase (ATGL), phosphorylated ATGL, perilipin and phosphorylated ACC in the scWAT or the vWAT between uninfected and infected animals nine weeks post-infection relative to the GAPDH loading control (Supp. Fig. 2F). Phosphorylated HSL and phosphorylated AKT were also similar in the muscle and liver of uninfected and infected mice relative to the GAPDH loading control (Supp. Fig. 2G). Inhibiting adipose triglyceride lipase (ATGL) has been shown to block wasting in murine cancer cachexia models (Das *et al*., 2011), however, attempts to block lipolysis whith the pharmacological inhibitor of ATGL atglistatin failed due to altered parasite growth (Nolan *et al*., 2017). Taken together, our data suggest that increased lipolysis was not the primary driver of chronic cachexia during *T. gondii* infection. This conclusion is consistent with the observed rebound in subcutaneous adipose weight and the stabilization of visceral weight adipose weight loss during chronic cachexia (Fig. 1E-F); as well as the clinical observations that not all patients present with adipose tissue loss and that cachexia can co-occur with obesity (Evans *et al*., 2008).

In the course of probing tissues by western blot we noticed that our β-actin loading control was significantly upregulated in infected mice relative to uninfected controls in all the tissues assessed (Supp. Fig. 2G-H). β-actin has over 90% sequence homology with activated fibroblast marker alpha smooth muscle actin (α-SMA). A recent report demonstrated that most commercially available antibodies raised against β-actin cross-react with α-SMA (Perrin and Ervasti, 2010). In response to tissue inflammation or mechanical stress, fibroblasts and other precursor cells trans-differentiate into myofibroblasts which are a major producer of extracellular matrix. This is an important aspect of wound healing but a dysregulated myofibroblast response can lead to fibrosis and impaired tissue function. (Nouchi *et al*., 1991; Carpino *et al*., 2005; Moreira, 2007). Based on this, we hypothesized that the increase in β-actin observed by immunoblots represented an increase in α-SMA and that metabolic tissues exposed to chronic inflammation during cachexia were becoming fibrotic. To test this directly, we assessed smooth muscle actin levels in tissue lysates from nine weeks post-infection or uninfected mice. Consistent with this hypothesis, smooth muscle actin was significantly upregulated in liver (Fig. 2A), muscle (Fig. 2B) and vWAT (Fig. 2C) at nine weeks post-infection relative to the GAPDH loading control. Tissues were sectioned and stained with Picrosirius Red, an anionic dye that forms sulfonic salt bridges with basic groups on elongated collagen fibers. Significant perivascular collagen deposition was observed in the liver (Fig. 2D), muscle (Fig. 2E) and vWAT (Fig. 2F) of infected mice relative to uninfected controls. Together these data are consistent with perivascular fibrosis in major metabolic tissues during chronic cachexia induced by *T. gondii* infection.

**Figure 2:**
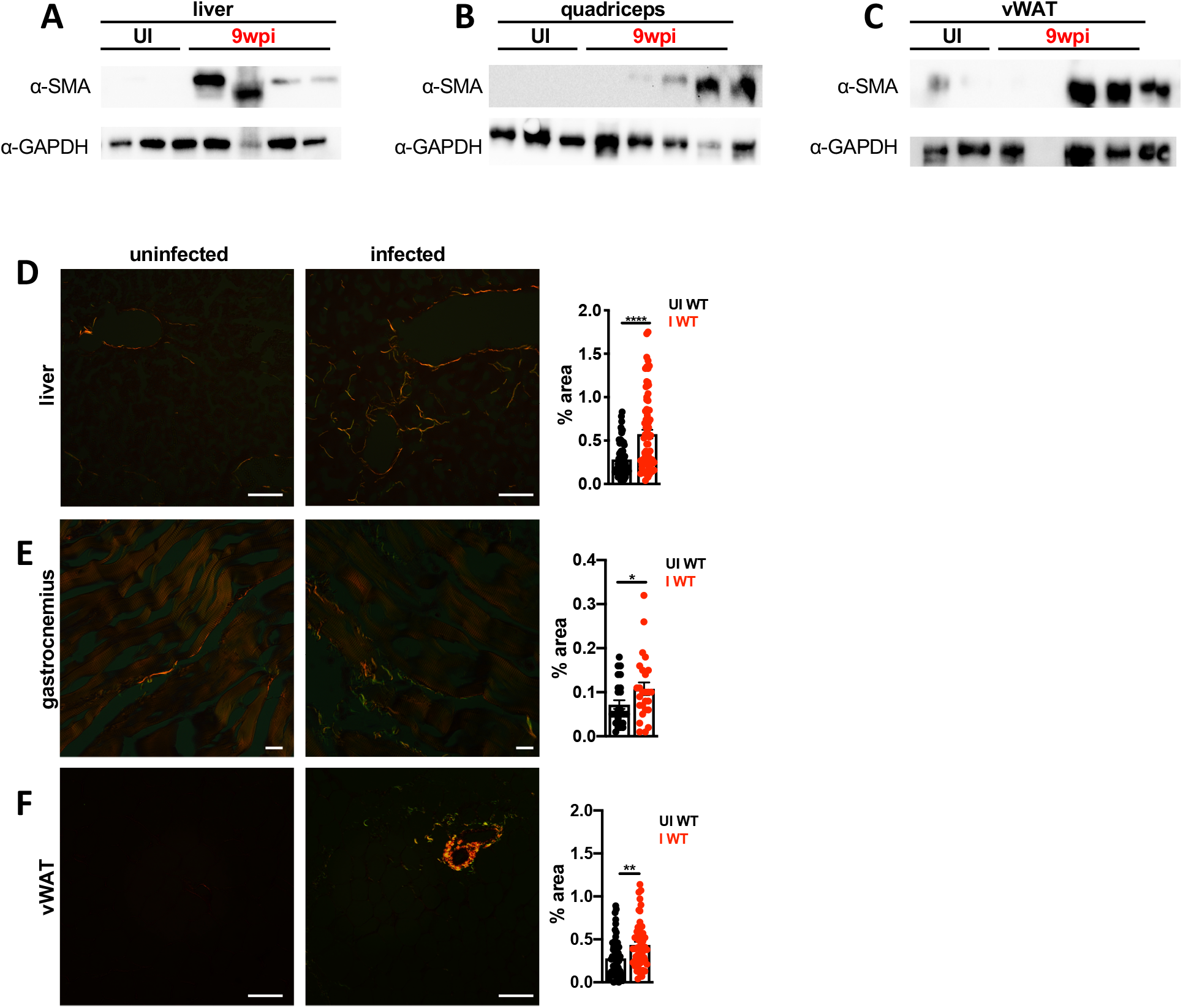
*Toxoplasma*-induced cachexia is associated with multi-organ fibrosis. **A-C** Western blot for alpha smooth muscle actin (α-SMA) or GAPDH on liver **(A)**, quadriceps **(B)**, and vWAT **(C)** tissue lysate at 9 wpi. Representative of at least 3 independent experiments. Each lane represents an individual mouse. **D-F**, Picrosirius red staining on formalin-fixed paraffin-embedded liver **(D)**, gastrocnemius **(E)**, or vWAT **(F)** using polarized light at 9 weeks post-infection. Representative images shown at left. Picrosirius red staining was quantified in ImageJ (right) and pixel density represented as % of each field of view. Each point represents a field of view (N=3 mice per group, 5-10 fields of view per mouse were quantified). Scale bars represent 50 μm. **D-E** are representative of at least 2 experiments and **(F)** is pooled between two independent experiments. Error bars are standard error of the mean. Statistical outliers were removed using the ROUT method (Q=1%), in **D-F.** *P < 0.05; **P < 0.01; ***P < 0.001 by unpaired Student’s T test.

### *T. gondii*-induced cachexia is dependent on IL-1 receptor signaling

TNF-α, IL-1, IL-6, and IFN-γ are inflammatory cytokines that are induced in both cachexia and *T. gondii* infection (de Matos-Neto *et al*., 2015). In *T. gondii* infection, mice deficient in the TNF-α, IL-6 or IFN-γ signaling pathways succumb to parasite overgrowth in acute or early chronic infection (Suzuki *et al*., 1988, 1996, 1997; Deckert-Schlüter *et al*., 1996; Jebbari *et al*., 1998; Yap and Sher, 1999; Schlüter *et al*., 2003). Although *T. gondii* can directly induce IL-1 release in vitro the role IL-1 plays during chronic *T. gondii* infection is comparatively understudied (Ewald, Chavarria-Smith and Boothroyd, 2014; Gorfu *et al*., 2014). This question was particularly interesting to us given the emerging role for the IL-1 axis in disease tolerance programs (Chang, Grau and Pechère, 1990; Alves-Rosa *et al*., 2002; Villeret *et al*., 2013; Bersudsky *et al*., 2014; Schieber *et al*., 2015; Rao *et al*., 2017).

To assess the role of the IL-1 axis in *T. gondii*-induced cachexia, mice deficient in the type I IL-1 receptor (IL-1R^-/-^) or control C57BL/6 (WT) mice were infected intraperitoneally with *T. gondii*. IL-1R^-/-^ mice had significantly improved long-term survival compared to WT mice (Fig. 3A). IL-1R^-/-^ and WT mice lost similar amounts of weight from 1-4 weeks post infection (Fig 3B), however, IL-1R^-/-^ mice regained weights by eleven weeks post-infection (Fig 3B). Importantly, the ability of IL-1R^-/-^ mice to rebound was not due to better parasite clearance as IL-1R^-/-^ mice had a similar chronic parasite load in the brain in terms of cyst number (Fig 3C, left), cyst size (Fig 3C, right, cysts can contain variable numbers of parasites) and by qPCR using primers specific to the *T. gondii* 529-bp repeat element (RE) to detect parasite genomic DNA (Fig. 3D) (Kasper *et al*., 2009). Infected IL-1R^-/-^ mice went through a period of anorexia comparable to the infected WT, but both genotypes regained eating to uninfected levels (Fig. 3E). IL-1 was originally named ‘anorexin;’ however, other inflammatory cytokines have subsequently been shown to induce anorexia so this result was not surprising (Plata-Salaman, 1998). Transient anorexia was consistent with the observation that both genotypes lost fat (Fig. 3F, left) and lean (Fig. 3F, right) body mass at two weeks post-infection by EchoMRI; however, IL-1R^-/-^ mice regained fat and lean body mass by nine weeks post-infection whereas infected WT mice did not (Fig. 3F). When individual tissues were measured, we found that IL-1R deficient mice lost significantly more muscle mass by two weeks infection than infected WT but regained muscle by nine weeks post infection (Fig. 3G, Quad, left) In contrast, infected WT mice continued to lose muscle mass by nine weeks post infection, which was significant relative to uninfected and IL-1R^-/-^ (Fig. 3G, Quad, left). Both WT and IL-1R^-/-^ mice regained subcutaneous adipose tissue; however, the vWAT remained significantly wasted at nine weeks post-infection (Fig. 3G). Both WT and IL-1R^-/-^ mice exhibit pronounced hepatomegaly two weeks post-infection. Unlike the infected wildtype mice, which progress to liver atrophy, the hepatomegaly resolved in IL-1R^-/-^ mice by nine weeks post-infection, and liver weights remained comparable to uninfected IL-1R^-/-^ mice (Fig. 3G).

**Figure 3:**
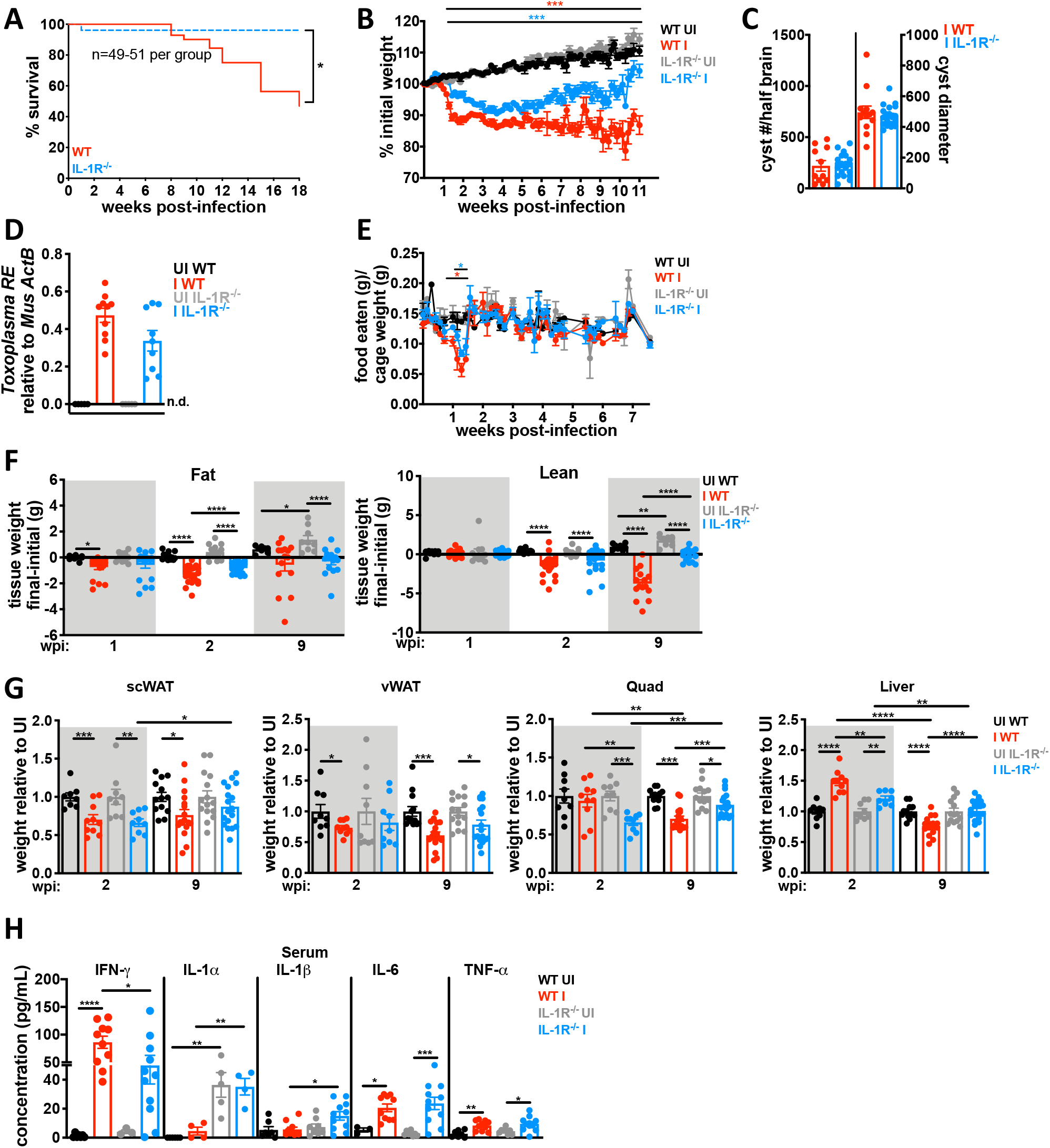
Acute cachectic weight loss is IL-1-independent, but IL-1R^-/-^ mice are protected from chronic *Toxoplasma*-induced cachexia. 12-14 week old C57Bl6 (WT, black UI, red I) or IL-1R^-/-^ mice (gray UI, blue I) were intraperitoneally infected with 10 *Toxoplasma* cysts of the Me49 background and monitored as described in Figure 1. **A**, Survival curves of infected WT or IL-1R^-/-^ mice N=49-51 mice per group, pooled from 8 independent experiments. p-value was determined by Log-rank test. **B**, Weight curves showing percent of initial weight. N=34-48 mice per group, pooled from 7 independent experiments. **C**, Brain cyst burden and diameter quantified at 9 wpi. N=11-21 mice per group, pooled from 3 independent experiments. **D**, qPCR for the *Toxoplasma* 529-bp repeat element in genomic DNA relative to mouse β-actin in brains 9 wpi. N= 5-10 mice per group, pooled from 2 independent experiments. **E**, Daily food intake per cage normalized to pooled weight of the mice in the cage measured every 24 hours. N=2-4 cages per group, pooled from 2 independent experiments. **F**, Echo MRI quantification of fat (left) and lean (right) tissue mass at 2 or 9 wpi. N=9-22 mice per group, pooled from 2-4 independent experiments. **G**, Inguinal subcutaneous white adipose tissue (scWAT), epididymal visceral white adipose tissue (vWAT), quadriceps (Quad) and liver weights shown relative to the mean weight of uninfected tissue at 2 wpi or 9 wpi. N=9-18 mice per group, pooled from at least 2 experiments. **H**, Serum cytokines measured by Luminex at 9 wpi. N=7-10 mice per group. IL-1α was measured by ELISA. N=4 mice per group. Error bars are standard error of the mean. *P < 0.05; **P < 0.01; ***P < 0.001, **** P < 0.0001 by unpaired Student’s T test with Holm-Sidak method to correct for multiple comparisons.

Importantly, levels of circulating IFN-γ, IL-6 and TNF-α remained elevated in the IL-1R^-/-^ mice nine weeks post-infection, consistent with the conclusion that these cytokines are not sufficient to sustain chronic cachexia during *T. gondii* infection (Fig. 3H). As expected based on their well-established roles in the restriction of parasite growth, sustained circulating IFN-γ, IL-6, and TNF-α were similar between genotypes (Fig. 3C-D). These data suggest that IL-1R signaling, which was dispensable to control parasite burden, plays a fundamentally different role in the biology of the immune response to *T. gondii* than the traditionally recognized pathogen targeting “resistance” mechanism(s) (Suzuki *et al*., 1988; Deckert-Schlüter *et al*., 1996; Jebbari *et al*., 1998; Schluter *et al*., 2003).

### IL-1 R signaling protects adipose and liver from acute pathology without affecting parasite load consistent with a role in disease tolerance

We next asked whether the protective phenotype in IL-1R^-/-^ metabolic tissues was established at acute infection. At two weeks post-infection, parasites are systemic and have infected most tissues in the mouse. Contrary to expectation, IL-1R^-/-^ had many large chalky lesions on the vWAT that were visible by eye (Fig. 4A, arrows). Although lesions were sometimes observed in the vWAT of infected WT mice, they were noticeably smaller and less frequent. These lesions resolved by nine weeks post-infection in both IL-1R^-/-^ and WT mice (Fig. 4A). Von Kossa staining of calcium deposits confirmed that lesions contained regions of necrosis (Fig. 4B, black) surrounded by inflammatory infiltrate (Fig. 4B, red nuclei)(Hanami, Hiraiwa and Yamamoto, 2012; Mineda *et al*., 2014; Misumi *et al*., 2019). H&E staining also showed that IL-1R^-/-^ vWAT had large, acellular regions surrounded by inflammatory infiltrate (Fig 4C). Although the composite histopathology score was similar between infected WT and IL-1R^-/-^, IL-1R^-/-^ vWAT had significantly more necrosis and at two weeks post infection than infected WT mice (Fig. 4D). The number of CD45^+^ immune cells (Supp. Fig. 3A) and the distribution of infiltrating immune cell populations (Supp. Fig. 3B) were similar between IL-1R^-/-^ and WT by flow cytometry. The levels of cachexia associated cytokines in vWAT lysate were similar by ELISA at two weeks post-infection (Supp. Fig 3C). These data indicate that the enhanced necrosis in the IL-1R^-/-^ vWAT was not simply due to a greater magnitude of inflammatory response in the absence of IL-1, although it is possible that more subtle differences in immune cell subpopulations exist. Importantly, parasite burden in the vWAT was similar between WT and IL-1R^-/-^ by qPCR, indicating that necrosis was also not due to parasite overgrowth in the absence of IL-1 signaling (Fig. 4E).

**Figure 4:**
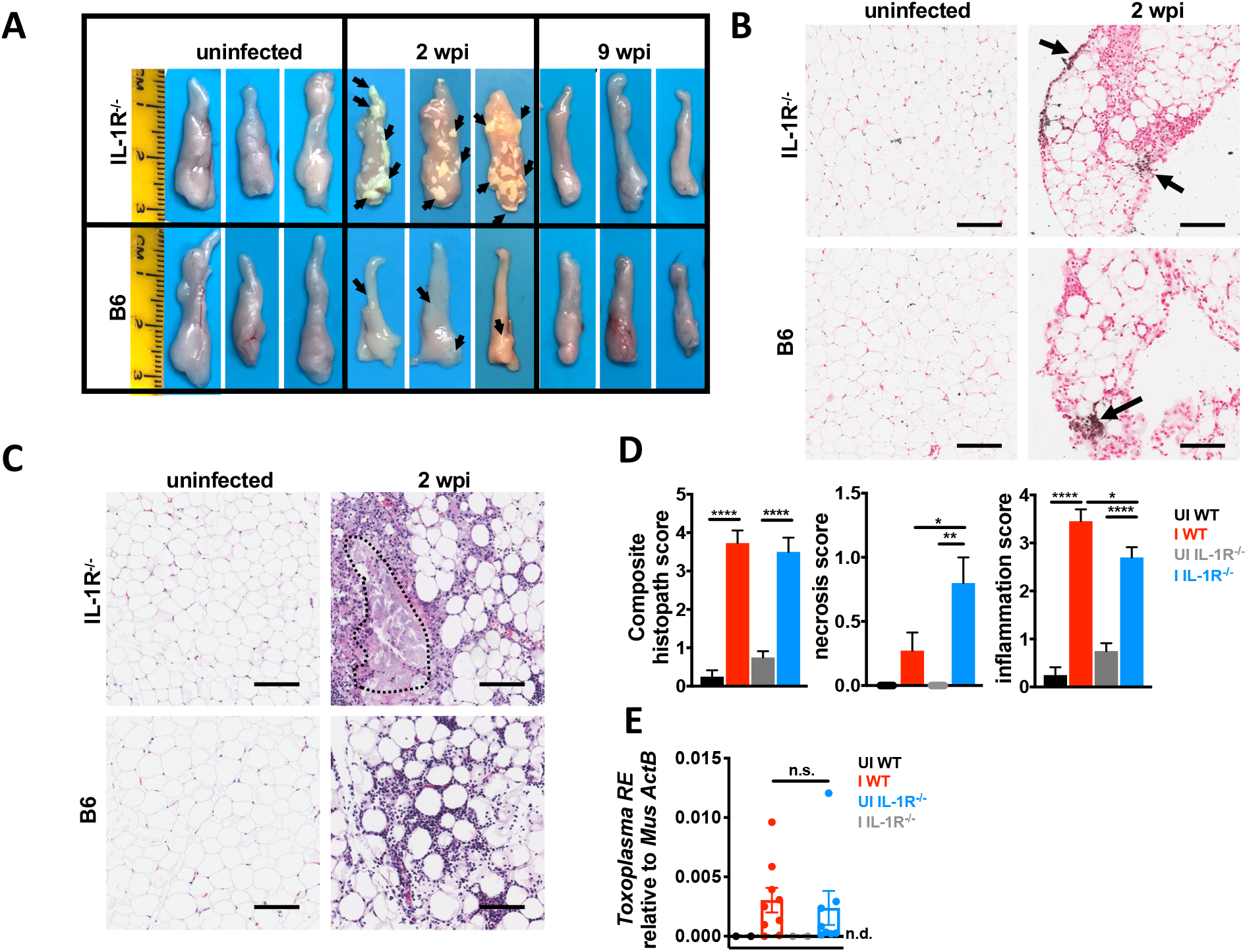
IL-1 confers protection from adipose tissue pathology during acute infection. 12-14 week old C57Bl6 (WT, black UI, red I) or IL-1R^-/-^ mice (gray UI, blue I) were intraperitoneally infected with 10 Me49 *Toxoplasma* cysts. **A**, vWAT was harvested from UI and I WT or IL-1R^-/-^ mice at 2 or 9 wpi. Representative of 5 independent experiments. **B**, Von Kossa staining on formalin-fixed paraffin-embedded vWAT at 2 wpi (n= 3 mice per group). Scale bar is 50 μm. **C-D**, Hematoxylin/eosin staining on formalin-fixed paraffin-embedded vWAT at 2 wpi. Scale bar is 50 μm. Necrotic lesion outlined in red. Histopathology quantified in D. N=6-8 mice pooled from 2 independent experiments. **E**, qPCR for the *Toxoplasma* 529-bp repeat element in genomic DNA relative to mouse β-actin in vWAT 2 wpi. N=2-9 mice per group, pooled from two independent experiments. Error bars are standard error of the mean, n.s.=not significant by unpaired Student’s T test with Holm-Sidak method to correct for multiple comparisons. *P < 0.05; **P < 0.01; ***P < 0.001, **** P < 0.0001 by unpaired Student’s T test with Holm-Sidak method to correct for multiple comparisons.

A similar observation was made in the liver at two weeks post-infection. WT and IL-1R^-/-^ livers had an equivalent number of focal perivascular immune infiltrate lesions by H&E staining (Fig. 5A-B), histopathology “composite” and “inflammation” scoring (Supp. Fig. 3G) as well as by flow cytometry for CD45^+^ cells (Supp. Fig. 3D). The distribution of infiltrating immune cells was similar between infected IL-1R^-/-^ and WT livers (Supp. Fig. 3E); although the number of CD4^+^ and CD8^+^ T cells in infected WT livers was not significant relative to uninfected due to mouse-to-mouse variability. IL-1R^-/-^ livers had significantly more enzymatically active caspase-3 staining in hepatocytes within inflammatory lesions than WT livers (Fig. 5C-D). Although the necrosis scores were not significantly different between WT and IL-1R^-/-^ by histopathology (Supp Fig. 3G), this may be due to better clearance of dying cells in the liver relative to the vWAT where lesions were much larger (Fig 4A, C). IL-1α, TNF-α and IFN-γ were significantly increased infected WT liver lysate compared to UI WT (Supp. Fig 3F). Cytokine levels trended similarly, but were more variable in infected IL-1R^-/-^ liver lysates (Supp. Fig. 3F). Parasite frequency is low in the liver and we were unable to confidently assess parasite load by qPCR, although parasite DNA was amplified above background in 1 of 11 of WT and 2 of 10 IL-1R^-/-^ livers using primers specific to the *T. gondii* RE or a *T. gondii* specific 35-fold repetitive gene, B1 (data not shown) (Burg *et al*., 1989; Jin *et al*., 2017).

**Figure 5:**
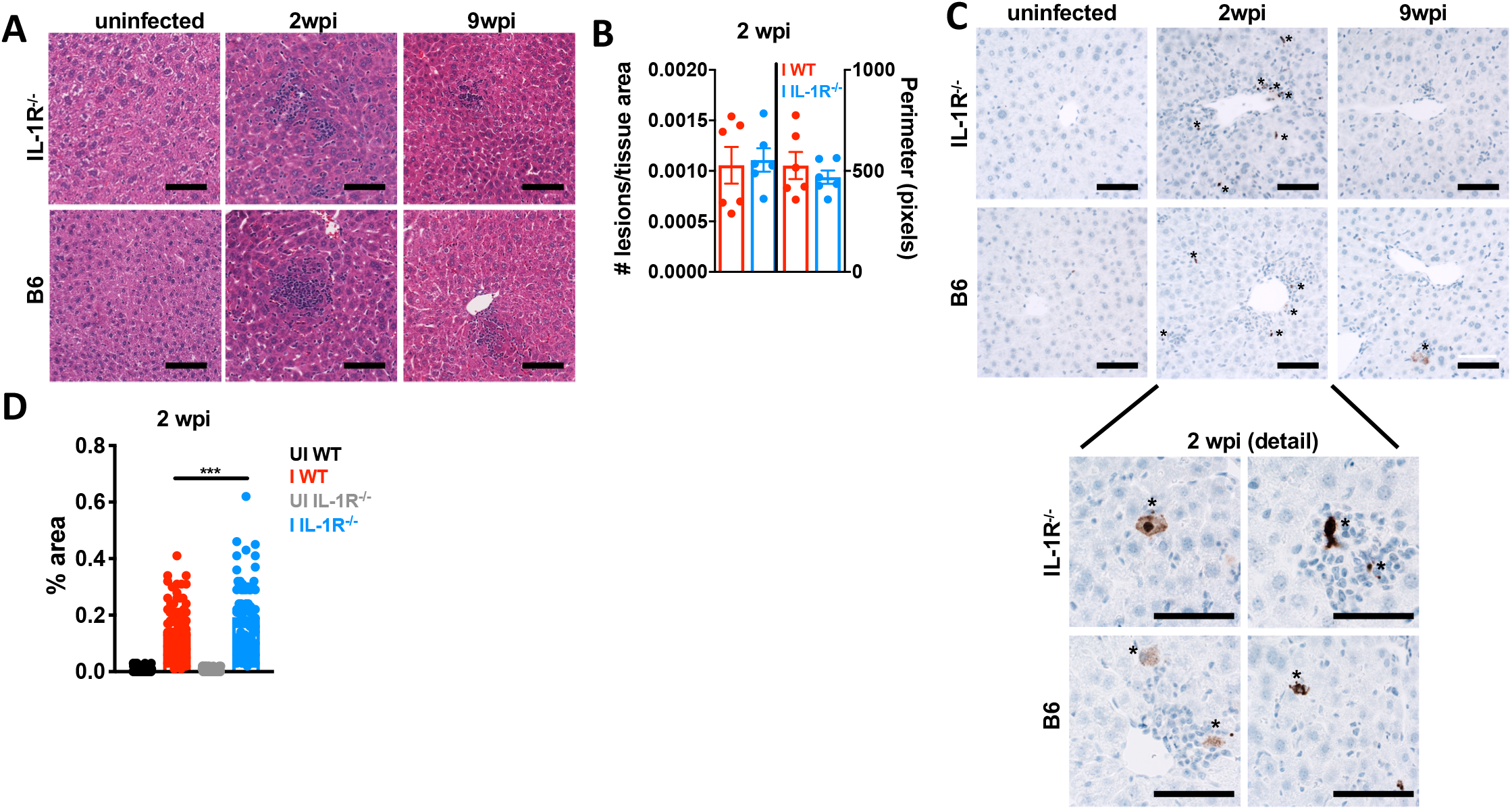
IL-1 signaling confers protection from liver tissue necrosis during acute infection. 12-14 week old C57Bl6 (WT, black UI, red I) or IL-1R^-/-^ mice (gray UI, blue I) were intraperitoneally infected with 10 Me49 *Toxoplasma* cysts. **A**, Hematoxylin/eosin staining on formalin-fixed paraffin-embedded liver at 2 and 9 wpi. **B**, Focal immune lesions were counted across slide scans of a whole liver section and lesion perimeter was measured using ImageJ. Each point represents an individual animal. N=3 mice per group, data pooled from 2 independent experiments. **C**, Immunohistochemistry for cleaved caspase-3 (brown positive staining marked by asterisks). Representative of 2 independent experiments. **D**, Cleaved caspase-3 staining area was quantified using ImageJ. Each point represents a blinded field of view. N=6 mice per group, 10 fields of view per mouse were quantified. Data pooled between two independent experiments. Scale bars represent 50 μm. Error bars are standard error of the mean. Statistical outliers were removed using the ROUT method (Q=1%), in **D.** *, P < 0.05; **, P < 0.01; ***, P < 0.001, by unpaired Student’s T test with Holm-Sidak method to correct for multiple comparisons.

Together, these data are consistent with a model where intact IL-1R signaling limits cell death in the fat and liver during acute *T. gondii* infection. In the fat, this protective effect is not due to better parasite clearance, consistent with an emerging role for IL-1 as a regulator of disease tolerance during acute infection. In the long run, however, chronic IL-1R signaling comes at the cost of cachexia.

### Cachectic mice have an IL-1R-dependent failure to recover from the acute extracellular remodeling response

We next asked whether the fibrosis observed in *T. gondii* infection-induced chronic cachexia was dependent on IL-1R signaling. At two weeks post-infection, both WT and IL-1R^-/-^ mice upregulated α-SMA relative to uninfected controls in the liver (Fig. 6A), vWAT (Fig. 6B) and muscle (Fig. 6C) relative to uninfected controls. Interestingly, in the vWAT which had the most robust acute pathology (Fig 4), upregulation of α-SMA was more pronounced in the IL-1R^-/-^ at two weeks post infection (Fig. 6B). At nine weeks post-infection WT mice exhibited sustained high levels of α-SMA in the liver and muscle and levels of α-SMA in the vWAT increased compared to uninfected mice and infected IL-1R^-/-^. By contrast, at nine weeks post-infection, IL-1R^-/-^ liver, vWAT and muscle had levels of α-SMA comparable to uninfected controls (Fig. 5A-C).

**Figure 6:**
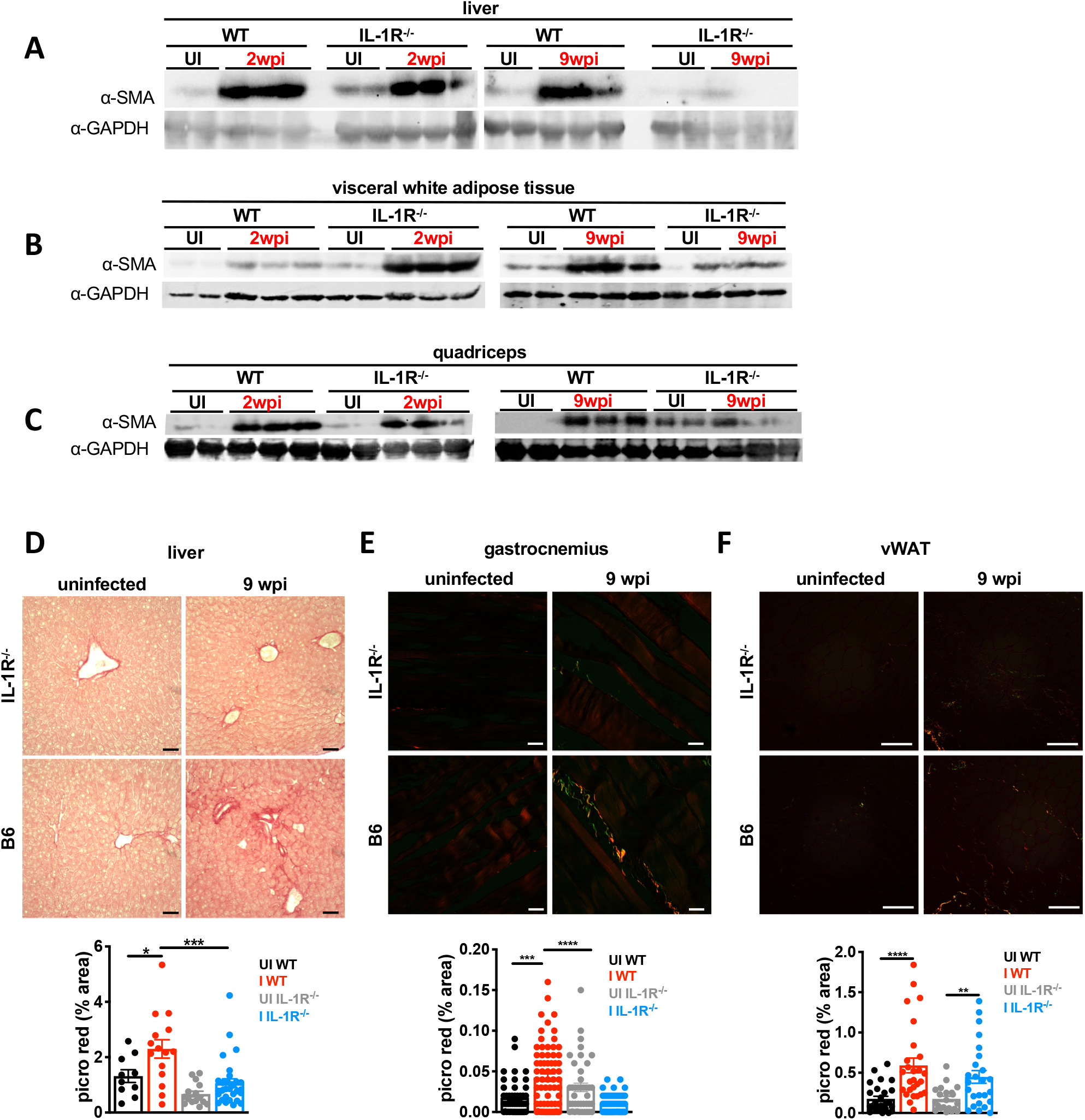
Cachectic mice develop IL-1-dependent liver and skeletal muscle fibrosis. **A-C** Western blot for alpha smooth muscle actin (α-SMA) or GAPDH on liver (A), vWAT (B), and quad (C) tissue lysate at 2 or 9 wpi. Representative of at least 2 independent experiments. Each lane represents an individual mouse. **D-F** Picrosirius red staining on formalin-fixed paraffin-embedded liver (D), gastrocnemius (E), or vWAT (F) using brightfield (D) or polarized light (E-F) at 9 weeks post-infection. Picrosirius red staining was quantified in ImageJ (below) and pixel density represented as % of each field of view. N=3 mice per group, 5-10 fields of view per mouse were quantified Representative of 2 experiments. Error bars are standard error of the mean. Scale bars represent 50 μm. Statistical outliers were removed using the ROUT method (Q=1%), in **E** and **F.** *P < 0.05; **P < 0.01; ***P < 0.001, ****P < 0.0001 by unpaired Student’s T test with Holm-Sidak method to correct for multiple comparisons.

To test whether α-SMA upregulation was associated with bona fide fibrosis, the vWAT, liver and muscle were sectioned and stained with Picrosirius Red. At nine weeks post-infection WT mice had more perivascular picrosirius red staining in the liver (Fig. 6D) and skeletal muscle (Fig. 6E) than infected IL-1R^-/-^ or uninfected mice. Colocalization of collagen I and collagen III staining with perivascular immune infiltrate and α-SMA positive fibroblasts was also assessed by immunofluorescence microscopy (Supp. Fig 4A-C). Even though α-SMA levels were reduced in the vWAT at 9 weeks post-infection (Fig 6C), Picrosirius Red staining was similar in the vWAT of infected WT and IL-1R^-/-^ (Fig. 6F). These data indicate that fibrosis in the vWAT is IL-1-independent or that the magnitude of necrosis in IL-1R^-/-^ vWAT at two weeks post-infection was too severe to recover from completely. These data support the conclusion that acute tissue remodeling initiated at two weeks post-infection is IL-1-independent; however, the IL-1 axis is necessary to sustain tissue remodeling in the muscle and liver resulting in perivascular fibrosis by nine weeks post infection.

### IL-1 expression is elevated in fibrotic tissues and IL-1 can directly trigger myofibroblast differentiation

We next asked whether IL-1 expression was sustained at local sites of fibrosis. At nine weeks post-infection, IL-1α and IFN-γ levels were significantly elevated in WT liver lysates relative to uninfected (Fig. 7A). In the muscle, cachexia-associated cytokines trended towards lower expression in infected WT relative to uninfected (Supp. Fig. 5A-B). In the liver, IL-1α positive cells were observed within collagen I positive inflammatory lesions; however, these cells did not co-stain with CD45, suggesting that they are not infiltrating immune cells (Fig. 7B). A caveat with these experiments is that IL-1 is synthesized and stored in the cells until it is released by cell death or damage indicating that the IL-1α we observed may not be biologically functional. However, infected WT mice had significantly elevated necrosis in the liver relative to uninfected mice at nine weeks post-infection, indicating a putative mechanism of release (Supp. Fig. 3H). IL-1R staining was also observed in proximity of collagen deposition on cells that co-stained for α-SMA (Fig. 7C). Cumulatively, these data are consistent with the conclusion that IL-1α is expressed in fibrotic regions of the liver that co-localize with α-SMA positive cells with fibroblast-like morphology expressing IL-1R.

**Figure 7:**
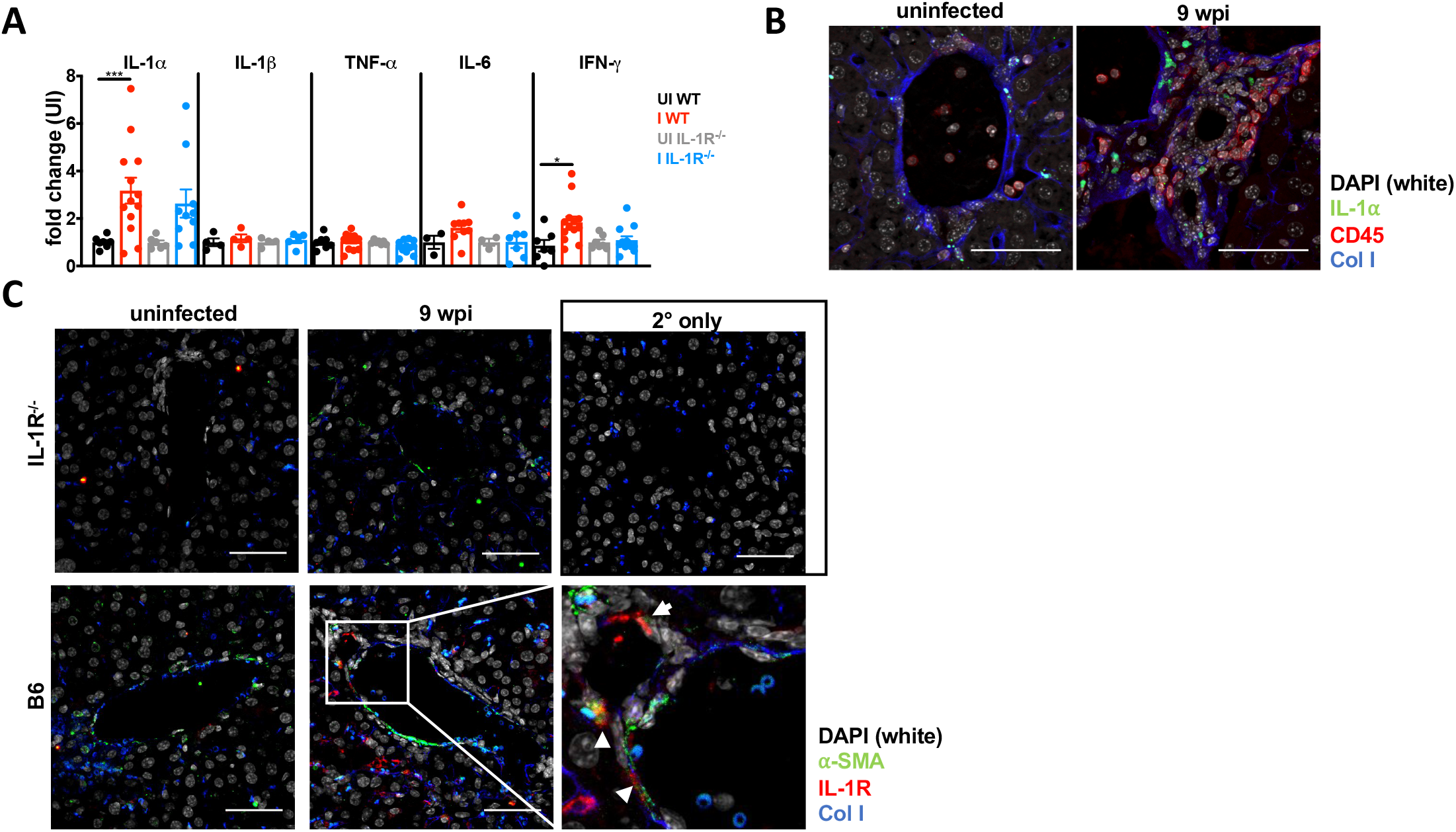
IL-1α and IL-1R are expressed in the liver at chronic infection. **A**, Cytokines in liver lysates from mice 9 wpi were measured by ELISA. Data are presented as fold change relative to the mean of uninfected levels. N=4-13 mice per group, pooled from two independent experiments. **B-C**, Immunofluorescence labeling of nuclei (DAPI white) IL-1α (green), CD45 (red) and collagen1α1 (blue) (**B**) nuclei (DAPI white) α-smooth muscle actin (green), IL-1R (red) and collagen1α (blue) in the liver of uninfected or 9 wpi WT or IL-1R^-/-^mice (**C**). Inset, arrow head represents α-smooth muscle actin, IL-1R co-staining cells (arrow heads). (**B-C**) represent maximum intensity projections of 9-13 μm thick z-stacks. Scale bar represents 50 μm. Error bars are standard error of the mean. *P < 0.05; **P < 0.01; ***P < 0.001 by unpaired Student’s T test with Holm-Sidak method to correct for multiple comparisons.

Based on the proximity between IL-1α, IL-1R expression and collagen deposition in the liver we next asked if IL-1 could promote activated fibroblast phenotypes directly. Murine embryonic fibroblasts (MEFs) were treated with media, IL-1α or TGF-β for 48 hours and then stained with phalloidin and an antibody specific to α-SMA for immunofluorescence imaging (Fig. 8A). Compared to the media control, IL-1α led to a significant increase in cell area (Fig. 8B), a measure of contractility associated with fibroblast activation, and α-SMA (Fig. 8C) to levels similar to treatment with TGF-β, the positive control. By contrast, neither IL-1α or IL-1β significantly impacted MEF cell proliferation or MEF cell survival under serum starvation conditions (Supp. Fig 5G).

**Figure 8:**
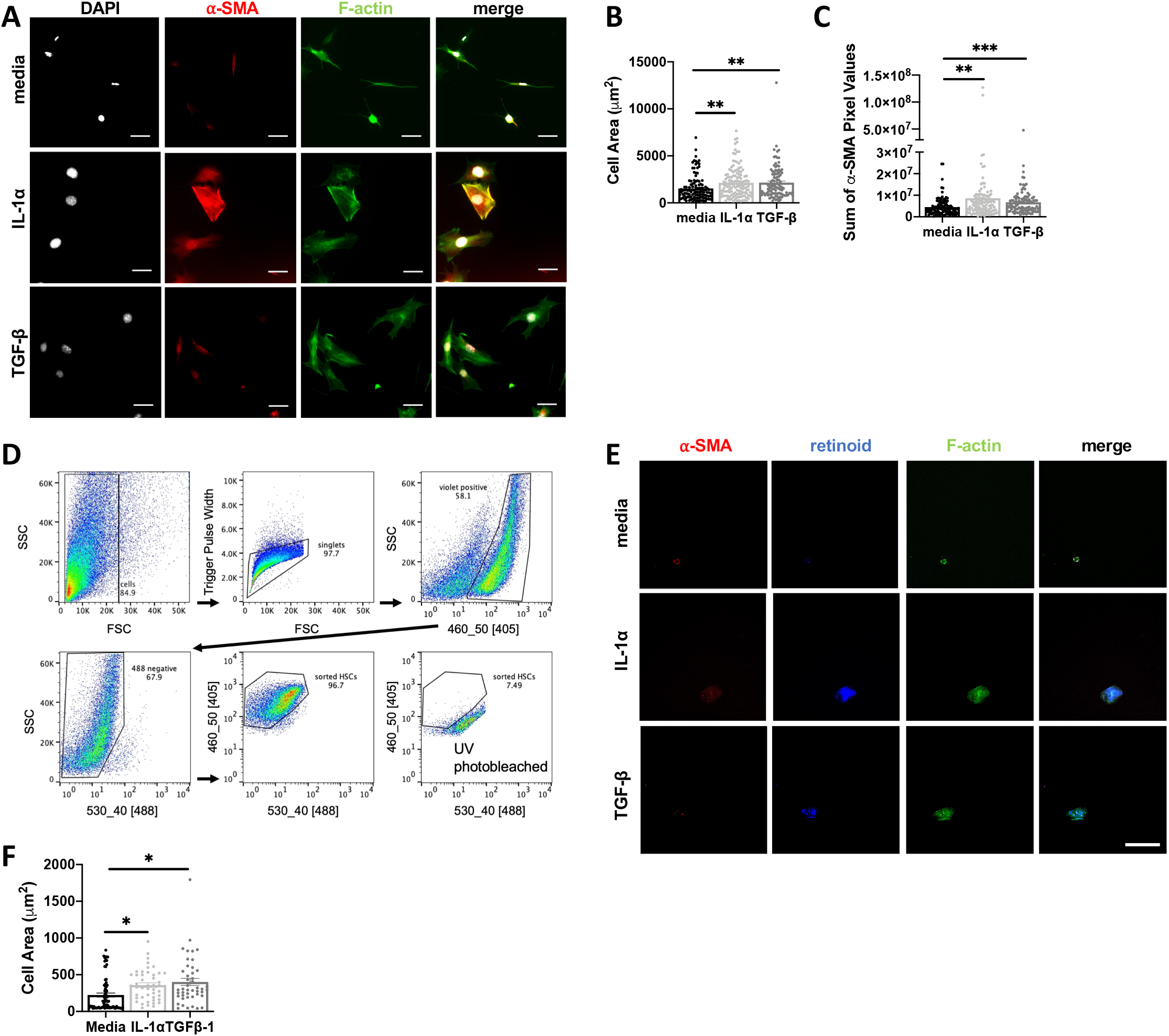
IL-1 can induce contractility in MEFs and primary hepatic stellate cells. **A-C**, MEF cells were incubated with media, 10ng/mL IL-1α or 5 ng/mL TGFβ-1 for 48 hours. After fixation, MEFs were stained for F-actin, alpha-smooth muscle actin (α-SMA), phalloidin and nuclei and cell spreading was quantified in **(B)**, and levels of α-SMA expression were quantified in terms of pixels/cell **(C). D-F**, Primary hepatic stellate cells (HSCs) were isolated from uninfected mouse livers, and FACS sorted based on endogenous retinoid fluorescence. Retinoid fluorescence was validated by UV photobleaching (**D**). **E-F**, HSCs were seeded onto 4kPa hydrogels coated with 10ug/mL of fibronectin and cultured with 10ng/mL IL-1α, 10 ng/mL TGF-β, or media alone for 48hrs and then fixed and stained for F-actin and α-SMA and imaged by confocal microscopy. Scale bar represents 50 μm. Total cell area quantified in (**F**) Error bars are standard error of the mean. *P < 0.05; **P < 0.01; ***P < 0.001, ****P < 0.0001 by unpaired Student’s T test.

To examine whether IL-1 was sufficient to induce myofibroblast differentiation in a liver-relevant cell type we took advantage of a robust protocol to isolate primary hepatic stellate cells (HSCs), the major myofibroblast precursor cell in the liver (Mederacke *et al*., 2015a). Following collagenase digestion, HSCs were FACS-sorted based on endogenous retinoid droplet autofluorescence (Mederacke *et al*., 2015b). Rigid structure mechanosensing can directly trigger myofibroblast differentiation (Pelham and Wang, 1997), potentially masking an effect of IL-1, so HSCs were plated on a 4kPA hydrogel to approximate the liver rigidity (Fig. 8D) (Yin *et al*., 2013). HSCs incubated with 10 ng/mL IL-1α or TGF-β for 48 hours significantly increased cell area, a measure of fibroblast contractility (Fig 8E, F). These data suggest that local IL-1R signaling in perivascular liver inflammatory lesions may directly trigger myofibroblast differentiation during chronic *T. gondii* infection-induced cachexia.

## Discussion

Here we show that intraperitoneal *T. gondii* infection is a robust model for chronic cachexia that recapitulates critical aspects of clinical cachexia. This is supported by several previous reports describing chronic weight loss and muscle dysfunction during *T. gondii* infection (Arsenijevic *et al*., 1997, 1998, 2001; Oldenhove *et al*., 2009; Hatter *et al*., 2018). Compared to other current models of cachexia a major advantage of the *T. gondii* infection model is its longevity, which opens the door to studying mechanisms of disease reversal and testing potential therapeutics (Konishi *et al*., 2015; Baazim *et al*., 2019). The duration of the *T. gondii* cachexia model allowed us to recognize that mice deficient in IL-1 signaling undergo acute cachectic weight loss similar to that observed in WT mice, but are protected from chronic cachexia. To our knowledge, this is the first example of a genetic rescue of cachexia after peak weight loss has occurred. This result is significant because it indicates that the inflammatory networks responsible for maintaining cachexia are distinct from those that initiate disease. Given the conserved role IL-1 plays across etiologies of clinical cachexia, we anticipate that the mechanisms uncovered in *T. gondii* cachexia may have important implications for cachexia in general (McDonald, McMillan and Laird, 2018).

One reason that the IL-1-dependent cachexia phenotype during *T. gondii* infection may have been missed until now is that the field has largely focused on immune mechanisms of parasite growth restriction and related host survival (so-called resistance mechanisms). However, the few reports looking at IL-1 signaling in *T. gondii* infection are consistent with the conclusion that IL-1 is not necessary to control parasite burden. In 2007 La Rosa and colleagues reported that IL-1R antagonist treatment did not affect parasite burden and stated that IL-1R^-/-^ mice are no more susceptible to *T. gondii* infection than wildtype controls (LaRosa *et al*., 2008). A 1998 study showed that IL-1α or IL-1β administration one day before infection protected Swiss Webster mice from an LD100 dose of type III *T. gondii* (Chang, Grau and Pechère, 1990). Although parasite load was not measured directly, all experimental groups had comparable *T. gondii* specific IgG and evidence of brain infection. In a more recent study IL-1R^-/-^ mice orally infected with Type II *T. gondii* had less ileitis and Paneth cell depletion than wildtype mice; and protection in the IL-1R^-/-^ mice corresponded with a reduced (rather than increased) parasite load (Villeret *et al*., 2013). Taken together, these data are consistent with an emerging role for IL-1 as a main regulator of tolerance biology, rather than pathogen restriction in the context of *T. gondii* infection (Rao *et al*., 2017; Benjamin *et al*., 2018).

“Disease tolerance mechanisms” are defined as shifts in homeostasis that improve host fitness by allowing tissues to function in the face of (or ‘tolerate’) damage caused by infection and inflammation without directly targeting the pathogen (Medzhitov, Schneider and Soares, 2012). Our results support a canonical disease tolerance role for the IL-1 signaling axis in acute *T. gondii* infection, because mice lacking IL-1R have more severe adipose tissue and liver pathology even though parasite load is similar to wildtype mice. Future experiments will be informative to understand how an intact IL-1 signaling axis is conferring protection to liver and adipose resident cells, and whether or not this is a direct effect. IL-1 has long been associated with anorexia, sickness behavior and metabolic shifts. In a recent study, *Salmonella typhimurium* mutants that could not inhibit IL-1β release in the intestine triggered a anorexic response via the afferent vagus nerve which ultimately lead to bacterial invasiveness and host death (Rao *et al*., 2017). However, in our model the gastrointestinal tract was bypassed, anorexia was similar between WT and IL-1R^-/-^ mice (Fig 3) and afferent vagotomy failed to rescue cachexia in WT mice (data not shown), suggesting a different mechanism is at play. Metabolic regulation at a cellular level has also been shown to be critical for modulating disease tolerance. Recently, T cell intrinsic type I interferon signaling was shown to regulate acute cachectic weight loss during LCMV infection. This response was CD8^+^ T cell dependent, however, viral burden was not measured directly in this study, so it is not clear if this is resistance or disease tolerance-mediated (Baazim *et al*., 2019). Another recent study has shown that cyclophilin D-dependent control of CD8^+^ T cells metabolism regulated disease tolerance in the lung during *Mycobacterium tuberculosis* infection, as cyclophilin D deficient CD8^+^ T cells were hyperproliferative, hyperactivated, and lead to lethal lung pathology without altering bacterial burden (Shen *et al*., 2017). Although a CD8^+^ T cell response is necessary to restrict *T. gondii* infection, Type I interferon is not robustly induced during disease due to the parasite secreted effector IST2 which blocks interferon response elements in infected cells (Gay *et al*., 2016). Moreover, a study from the Hunter lab showed that IL-1R signaling on T cells had no effect on the progression of *Toxoplasma* infection(LaRosa *et al*., 2008). We did not observe any significant difference in liver or adipose tissue T cell recruitment (Supp. Fig. 3B, E, Supp. Fig. 5D, F) or activation state (data not shown) Future work will be necessary to determine whether T cell intrinsic mechanisms control cachexia progression is induced by IL-1 to control disease tolerance to *T. gondii* or a fundamentally distinct mechanism is at play.

Understanding how to target and promote tolerance biology has become a topic of clinical interest because such a strategy has the potential to mitigate the negative side effects of inflammation without putting the patient at increased risk of infection. However, an extension of the tolerance/resistance paradigm is that disease tolerance programs must come at a cost to the organism otherwise they would be selected for homeostatic use. Consistent with this theoretical framework, mice with an intact IL-1 signaling axis are protected from acute cachexia pathology, but ultimately fail to return to homeostasis. Although serum IL-1α and IL-1β are not significantly elevated in chronic infection, IL-1α is elevated in liver and possibly other tissue microenvironments. These data are consistent with a model in which cachexia can be viewed as a cost of long-term reliance on IL-1 mediated tolerance biology.

From the perspective of advancing our understanding of cachexia biology, *T. gondii* infection is a unique tool to uncover the role inflammatory mediators play in cachexia because the parameter of pathogen growth can be used to distinguish between immune effectors primarily involved in resistance versus tolerance biology. IL-1, IL-6, TNF-α and IFN-γ are an inflammatory signature of cachexia and *T. gondii* infection (de Matos-Neto *et al*., 2015; Hatter *et al*., 2018). IL-6, TNF-α and IFN-γ are absolutely critical for *T. gondii* resistance: mice deficient in these cytokines, their receptors, or downstream signaling cascades die from parasite overgrowth in acute infection or early chronic infection (Oliff *et al*., 1967; Fong *et al*., 1989; Matthys *et al*., 1991; Plata-Salaman *et al*., 1997; Haddad *et al*., 2005). Of note, TNF-α targeting therapeutics have failed to treat clinical cachexia. Interestingly, a first in class monoclonal antibody targeting IL-1α (MABp1) was recently tested in a phase 1 dose escalation trial for metastatic cancers. 70% percent of MABp1-treated patients exhibited increased lean body mass, and MABp1 significantly reduced sera IL-6 and increased global quality of life scores (Hong *et al*., 2014). Although cachexia was not an entry criteria for this study, 77% of the participants were actively losing weight in the six months leading up to the trial. Blocking resistance effectors like TNF-α should decrease damage to self, but may not correct homeostatic shifts that occur in response to the inflammatory environment. It is possible that targeting tolerance effectors like IL-1 directly, or in tandem with resistance mechanism inhibitors, may be a more efficacious way to restore homeostasis during cachexia. Future clinical trials with MABp1 in disease matched cachectic and weight stable patients will be informative.

What is the significance of IL-1-mediated tolerance on host fitness? Compared to cachectic WT mice, IL-1R^-/-^ have better survival during chronic infection, while wildtype mice have increased mortality, impaired activity, and reduced muscle mass. These findings are consistent with a 2018 study from the Wohlfert lab which found that chronically infected mice had muscle inflammation and impaired muscle strength supported by local T regulatory cells (Oldenhove *et al*., 2009). These phenotypes may translate to the debilitating co-morbidities of pain and fatigue that negatively impact the cachectic patient’s quality of life and ability to survive physically taxing treatments or secondary infection. For example, a study by Arsenijevic and colleagues demonstrated that the *T. gondii* infected mice with the most severe chronic weight loss were more susceptible to sub-lethal doses of endotoxin injection (Arsenijevic *et al*., 1998).

The mechanism(s) by which IL-1 is controlling chronic cachexia in *T. gondii* infection remains to be addressed. Mice deficient in the IL-1R antagonist have reduced fat mass and increased energy which prompted us to first investigate shifts in metabolic homeostasis as drivers of *T. gondii* infection-induced cachexia (Matsuki *et al*., 2003; Burke *et al*., 2018). However, we did not observe major increases in lipolysis or markers of non-shivering thermogenesis in our experiments. The most robust IL-1 dependent phenotype in cachexia was perivascular fibrosis in the muscle and liver of cachectic mice. This observation is exciting because adipose fibrosis has been observed in cachectic gastrointestinal cancer patients relative to age and disease matched, weight stable cancer patients (Batista *et al*., 2016; Alves *et al*., 2017). Our experiments showed that acute tissue remodeling is IL-1-independent. However, IL-1R deficient mice, which rebound their lean body mass and are protected from liver atrophy, are also protected from developing perivascular fibrosis in the liver and muscle. This finding is consistent with several reports that IL-1R1 deficient mice or mice treated with IL-1R antagonists are protected from liver fibrosis induced by bile duct ligation, or thioacetamide, as well as fibrosis induced by alcoholic liver injury (Gieling, Wallace and Han, 2009; Petrasek *et al*., 2011; Ganz *et al*., 2015; Meier *et al*., 2019). Mice deficient in IL-1α or IL-1β are protected from the transition between steatosis and liver fibrosis induced by hypercholesterolemia and acetaminophen induced liver injury (Kamari *et al*., 2011; Zhang *et al*., 2018). Fibroblast specific deletion of IL-1 also reduced fibrotic tissue remodeling after cardiac infarction (Bageghni, Drinkhill and Turner, 2019). These data suggest that IL-1 signaling is necessary for fibrosis in a range of tissue settings, including the liver and muscle. While IL-1 polymorphisms in humans haven’t been extensively studied in the context of cachexia, they have been associated with increased risk of fibrotic and liver disease (Takamatsu *et al*., 1998; Whyte *et al*., 2000; Hutyrová *et al*., 2002; Barlo *et al*., 2011; Alcaraz-Quiles *et al*., 2017; Tak *et al*., 2018). Our data indicate that the perivascular fibrosis associated with chronic cachexia during *T. gondii* infection is also IL-1 dependent.

Our data are consistent with the hypothesis that IL-1R signaling may directly influence fibroblast biology in *T.gondii*-induced cachexia. In the liver, IL-1α and IL-1R expressing cell types are observed within perivascular fibrosis lesions. Moreover, hepatic stellate cells, which are the major myofibroblast precursor in the liver are protected from cell death in direct response to IL-1 (Pradere *et al*., 2013). However, fibroblasts are extremely heterogenous in the liver and across tissues. Future studies will be necessary to identify precise myofibroblast cell or precursor populations that express IL-1R and test whether selectively eliminating or driving IL-1R signaling on these cell types is sufficient for fibrosis in the *T. gondii* infection.

Additional experiments will also be necessary to determine if fibrosis as causal, symptomatic, or preventing recovery from cachexia. Like cachexia, fibrosis has been notoriously difficult to target and reverse in the clinic. Recently, HDAC inhibitors have emerged as tools to halt fibrosis in the setting of cancer, as well as cardiac and pulmonary fibrosis (Kim *et al*., 2018; Lyu and Sanders, 2019). However, we found that the HDAC inhibitors losartan and scriptaid altered *T. gondii* growth in vivo and caused death of treated mice, confounding the interpretation of these experiments (data not shown). If fibrosis is necessary for cachexia progression, our data suggest that liver and/or muscle fibrosis are likely culprits because IL-1R^-/-^ mice were protected from cachexia but still exhibited visceral adipose fibrosis. There have only been a handful of studies assessing fibrosis in cachectic patients (Batista *et al*., 2016; Alves *et al*., 2017); however, there is abundant literature associating cachexia with fibrotic diseases. Cachexia is extraordinarily common in all etiologies of chronic liver disease. Emerging clinical data show a strong association between muscle wasting, disease progression, and mortality in all etiologies of chronic liver disease and in hepatocellular carcinoma (Loza *et al*., 2012; Hayashi *et al*., 2013; Ishimoto *et al*., 2013; D *et al*., 2015; Kalafateli *et al*., 2015; Dasarathy and Merli, 2016; van Vugt *et al*., 2016; Watt *et al*., 2017; Koo *et al*., 2017; Ukwaja *et al*., 2017; Han *et al*., 2018). In chronic obstructive pulmonary disease, which is driven by progressive lung fibrosis, and chronic heart disease cachexia rates range from 5-20% (von Haehling and Anker, 2010, 2014). Assessing fibrosis in cachectic patients is difficult because tissues like liver are not usually biopsied; however, muscle biopsy and assessment of adipose tissue removed during surgery are becoming increasingly common. Future experiments exploring the fibrotic landscape of cachectic patients will be informative. To date, these observations are largely correlative, but suggest that further investigation into the relationship between fibrosis and cachexia are warranted.

The major finding of our studies is that IL-1R deficient mice are protected from chronic cachexia and the associated muscle and liver fibrosis. The causal relationship between these diseases is an exciting new area for exploration in both animal models and clinical cachexia. Our data suggest that targeting signaling cascades that regulate disease tolerance biology, potentially in tandem with approaches to block pro-inflammatory resistance cascades, may be a new approach to identifying to developing treatments for chronic diseases like cachexia and fibrosis.

## Materials and Methods

### Infections

To generate cysts, 8-10 week female CBA/J mice were infected with 3-10 Me49 or Me49 stably expressing green fluorescent protein and luciferase (Me49-GFP-luciferase) bradyzoite cysts by intraperitoneal injection. 4–8 weeks following infection, mice were euthanized with CO_2_ and brains were harvested, homogenized through a 70 μm filter, washed 3 times in PBS, stained with dolichos biflorus agglutinin conjugated to either FITC or rhodamine(Vector labs) and the number of cysts were determined by counting FITC-positive cysts at 20x magnification using an EVOS FL imaging system (Thermo Fisher).

10-14 week male mice were infected with 10 Me49 or Me49-GFP-luciferase bradyzoite cysts by intraperitoneal infection. Prior to infection, mice were cross-housed on dirty bedding for two weeks to normalize commensal microbiota.

### Mouse strains/husbandry

C57BL/6 and IL-1R^-/-^ mice were purchased from Jackson Laboratories. All mice were bred in-house. All animal protocols were approved by the University of Virginia Institutional Animal Care and Use Committee (protocol # 4107-12-18). All animals were housed and treated in accordance with AAALAC and IACUC guidelines at the University of Virginia Veterinary Service Center.

### Assessment of parasite burden

To measure parasite burden, brains were harvested and homogenized through a 70 μm filter, washed 3 times in PBS, stained with dolichos biflorus agglutinin-FITC or -rhodamine (Vector labs) and the number of cysts were determined by counting biflorus agglutinin positive cysts at 10x magnification on an EVOS FL microscope (Life Technologies). Cyst diameter was quantified with ImageJ.

To measure parasite burden in the brain by qPCR, brains were harvested, homogenized in TRIzol Reagent (Invitrogen). Following chloroform extraction and removal of the upper aqueous layer and interphase, DNA was isolated by ethanol precipitation, washed in twice in 0.1 M sodium citrate in 10% ethanol, resuspended in 8 mM sodium hydroxide and quantified by NanoDrop One^c^ (Thermo Scientific).

30 ng brain genomic DNA was used for qPCR analysis of parasite burden. Primers and probes designed against the 529bp Repeat Element (RE) were obtained from IDT (*Forward:* 5’-CACAGAAGGGACAGAAGTCGAA-3’, *Reverse:* 5’-CAGTCCTGATATCTCTCCTCCAAGA-3’, *Taqman Probe:* 5’-CTACAGACGCGATGCC-3’). Quantity of parasite DNA are presented relative to host β-actin (Mm02619580_g, ThermoFisher Scientific). Isolation of DNA from adipose tissue for parasite burden analysis was performed using NucleoSpin DNA Lipid Tissue kit (Machery-Nagel) and following the manufacturer’s instructions. 10 ng adipose tissue genomic DNA were used for qPCR analysis. Data were analyzed using the ΔCt method (relative to host *actβ* as a reference gene).

### Tissue Weights and Body Mass Composition

Following euthanasia by CO_2_ and exsanguination, tissues were excised and frozen on dry ice in preweighed tubes. Tubes were then weighed to determine tissue mass. For tissues with bilateral symmetry (scWAT, vWAT, quad), both sides were excised, weighed, and averaged. Body mass composition was determined by using the EchoMRI™-100H Body Composition Analyzer.

### Histology

Following euthanasia and harvest, tissues were fixed overnight in 4% paraformaldehyde at 4°C, after which they were submitted to the Research Histology Core at the University of Virginia for paraffin-embedding, sectioning, and staining with either hematoxylin and eosin, or Picrosirius red. Von Kossa staining was performed using a Von Kossa Stain Kit (American Mastertech Scientific, Inc, Lodi, CA).

### Picrosirius red imaging

Slides were imaged on an Olympus BX51 microscope with an Infinity 1 camera (Lumenera) or a Zeiss Apotome2 (Carl Zeiss, Germany) under polarized light using the 20X or 40X objective. 5-10 blinded fields of view were acquired per mouse. To quantify percent area, images were binarized in Fiji (Schindelin *et al*., 2012), thresholded, and percentage of positive pixels per area was determined.

### Immunohistochemistry

Formalin-fixed paraffin-embedded tissue sections were stained for cleaved caspase-3 by the Biorepository and Tissue Research Facility at the University of Virginia. 5-10 blinded fields of view were acquired for each mouse using a 20x objective on an Olympus BH-2 microscope with an Infinity 1 Lumenera camera. To quantify percent area, using the color deconvolution plug-in (Ruifrok and Johnston, 2001), color channels were separated, thresholded according to a negative control, and percent of positively stained pixels was measured.

### Pathology Scoring

Liver immune lesions were counted across whole slide scans of H&E stained tissue, and perimeter was measured using Fiji. A blinded pathologist scored H&E slides using the following rubric: For the adipose tissue: Inflammation was scored as follows: 0) no lesions, 1) 1-10% scattered individual immune cells, 2) small aggregates and/or perivascular cuffs, 3) larger but still individual nodules, 4) coalescing nodules or perivascular accumulations of immune cells. Necrosis was scored as follows: 0) 0% of section affected, 1) 1%-25%, 2) 26%-50%, 3) 51%-75%, 4) 76%-100%. For the liver tissue, inflammation and EMH were scored as follows: 0) 0-5% of section affected, 1) 6%-25%, 2) 26%-50%, 3) 51%-75%, 4) 76%-100%. Necrosis was scored the same as in the adipose tissue.

### Immunofluorescence

Following euthanasia and harvest, tissues were fixed overnight in 4% paraformaldehyde at 4°C, after which they were submerged in 30% sucrose in PBS, embedded in Tissue-Tek^®^ O.C.T. Compound (VWR) and flash frozen on dry ice. Samples were submitted to the Research Histology Core at the University of Virginia for sectioning.

For IL-1α staining, slides were blocked for 1 hour at room temperature in blocking buffer (2% donkey serum, 2% goat serum, 0.1% Triton-X, 0.05% Tween-20). Samples were incubated overnight at 4°C in primary antibody diluted in blocking buffer (R&D AF-400-NA goat anti-IL-1 α, 1:50; Novus Biologicals NB600-408 rabbit anti-collagen I, 1:50; Biolegend 103101 rat anti-CD45, 1:50). The next morning, samples were washed 3 times in PBS/0.1% Triton-X and were incubated in secondary antibody (donkey anti-goat Dylight 594, Novus NBP1-75607, 1:500; Invitrogen A21245 goat antirabbit AF647, 1:300; and donkey anti-rat AF488, 1:500), for 1 hour at room temperature, diluted in blocking buffer. Samples were washed 3 times in PBS, stained for 5 minutes in 10 μg/mL DAPI, washed 3 times in PBS, and mounted in Vectashield. Slides were imaged on a Zeiss LSM 880 confocal microscope using the 40X or 63X objectives.

For IL-1R staining, antigen retrieval was performed by boiling samples in sodium citrate buffer (10 mM sodium citrate buffer + 0.05% Tween-20, pH 6.0) for 20 minutes and then washing 3 times in PBS. They were blocked for 1 hour at room temperature in 2% donkey serum and CD16/CD32 (Biolegend, Clone 93, 1:200). Samples were incubated overnight at 4°C in primary antibody diluted in blocking buffer (R&D AF771 goat anti-IL-1R, 1:50; Novus Biologicals NB600-408 rabbit anticollagen 1, 1:50). The next morning, samples were washed 3 times in PBS/0.1% Triton-X and were incubated in secondary antibody (Novus NBP1-75607 donkey anti-goat Dylight 594, 1:250 and LifeTech A21206 donkey anti-rabbit AF488, 1:200), and the directly conjugated α-SMA primary (Novus Biologicals NBP2-34760APC, 1:400) for 1 hour at room temperature, diluted in PBS/0.1% Triton-X. Samples were washed 3 times in PBS, stained for 5 minutes in 10 μg/mL DAPI, washed 3 times in PBS, and mounted in Vectashield (Vector Laboratories).

For collagen I and III staining, samples were boiled in sodium citrate buffer (10 mM sodium citrate buffer + 0.05% Tween-20, pH 6.0) for 20 minutes and then washed 3 times in PBS + 0.1% Triton-X. They were blocked for 1 hour at room temperature in blocking buffer (2% donkey serum + 2% goat serum)

Samples were incubated overnight at 4°C in primary antibody diluted in blocking buffer (Biolegend 103101 rat anti-CD45, 1:50; Novus Biologicals NB600-408 rabbit anti-collagen I, 1:50; Novus Biologicals NB600-594 rabbit anti-collagen III, 1:50). The next morning, samples were washed 3 times in PBS + 0.1% Triton-X and were incubated in secondary antibody (Invitrogen A-11007 goat-anti rat AF594, 1:500; LifeTech A21206 donkey anti-rabbit AF488, 1:200), and the directly conjugated α-SMA primary (Novus Biologicals NBP2-34760APC, 1:400) diluted in PBS + 0.1% Triton-X for 1 hour at room temperature. Samples were washed 3 times in PBS, stained for 5 minutes in 10 μg/mL DAPI, washed 3 times in PBS, and mounted in Vectashield (Vector Laboratories). Slides were imaged on a Zeiss LSM 880 confocal microscope (Carl Zeiss) using the 40X oil (numerical aperture: 0.09) or 63X oil (numerical aperture: 0.09) objectives and ZenBlack software (Carl Zeiss).

### Cytokine Measurements

Sera cytokines were measured by Luminex at the University of Virginia Flow Cytometry Core, or by ELISA (Thermo Fisher Scientific). Cytokine measurement on tissue homogenate was assessed using ELISA (Mouse IFN gamma Uncoated ELISA (Invitrogen, #88-7314); Mouse TNF alpha Uncoated ELISA (Invitrogen, #88-7324); IL-1 alpha Mouse Uncoated ELISA Kit (ThermoFisher #88-5019-88); eBioscience Mouse IL-6 ELISA Ready-SET-Go! Kit (Fisher Scientific #50-112-8863; IL-1 beta Mouse Uncoated ELISA Kit (Thermo Fisher Scientific #88-7013-88).

Sera cytokines were measured by Luminex at the University of Virginia Flow Cytometry Core, or by ELISA (Thermo Fisher Scientific). Cytokine measurement on tissue homogenate was assessed using ELISA kits according to the manufacturer instructions (Mouse IFN gamma Uncoated ELISA, Invitrogen, #88-7314; Mouse TNF alpha Uncoated ELISA, Invitrogen, #88-7324; IL-1 alpha Mouse Uncoated ELISA Kit, ThermoFisher #88-5019-88; eBioscience Mouse IL-6 ELISA Ready-SET-Go! Kit, Fisher Scientific #50-112-8863; IL-1 beta Mouse Uncoated ELISA Kit, Thermo Fisher Scientific #88-7013-88).

### Sorting and culturing of primary hepatic stellate cells

Primary murine hepatic stellate cells were isolated as previously described.(Mederacke *et al*., 2015a) Following sorting on a BD Influx Cell Sorter in the University of Virginia Flow Cytometry Core, cells were seeded onto fibronectin coated 4 kPa polyacrylamide hydrogels (Matrigen) and stimulated with 10 ng/mL recombinant mouse IL-1α (R&D), 10 ng/mL recombinant mouse TGF-β (R&D), or media alone for 48 hours, after which hydrogels were fixed with 4% paraformaldehyde and stained with phalloidin-488 (Invitrogen) and anti α-SMA antibody (Invitrogen, clone IA4). Cells were mounted with ProLong Diamond Anti-Fade mountant (Thermo Fisher Scientific), and imaged at room temperature on a Nikon Eclipse Ti microscope with an UltraView VoX imaging system (PerkinElmer) using a Nikon N Apo LWD 40X water objective (numerical aperture: 1.15) and cell area and α-SMA intensity were determined using Volocity software.

### Western blots

Flash-frozen tissues were homogenized by bead-beating in lysis buffer (25 mM Tris-HCl, 15 mM NaCl, 1 mM MgCl2, 2.7 mM KCl, 1 mM EDTA, 1 mM EGTA, 1 mM DTT, 1% Triton-X, and protease and phosphatase inhibitors (Roche EDTA-free protease inhibitor mini and Pierce™ Phosphatase Inhibitor Mini Tablets), and centrifuged. Protein concentration was measured using via BCA concentration (Pierce BCA Protein Assay Kit, Cat# 23225). Protein concentrations were equalized by diluting with 2X Laemmli buffer and lysate was then boiled at 95°C for 3 minutes. All protein samples were separated on 10% bis-Tris gels by SDS-PAGE and then transferred to PVDF membranes membranes using Trans-Blot Turbo Transfer System (Biorad). The membranes were blocked in 2% ECL Prime Blocking Reagent (GE Amersham RPN418) in TBS-T for 30 min, incubated in primary antibody for 1 hour at room temperature, washed 3X in TBS-T, incubated in secondary antibody for 30 min at room temperature, washed 3X in TBS-T and then imaged on either the Bio-Rad ChemiDoc Imager (for HRP secondary antibodies) or the Leica Typhoon (for fluorescently-conjugated secondary antibodies). The following primary antibodies were used to probe protein levels by immunoblotting: anti-GAPDH (Cell Signaling Technology, clone D16H11), anti-β-Actin (Santa Cruz Biotechnology, clone 1), anti-α-SMA (Santa Cruz Biotechnology, clone CGA7), anti-pACCα (Santa Cruz Biotechnology, clone F-2), anti-ATGL (Santa Cruz Biotechnology, clone F-7), anti-HSL (Cell Signaling Technology, #4107), anti-phospho-HSL (Ser660) (Cell Signaling Technology, #4126), anti-AKT (Cell Signaling Technology, clone C67E7), and anti-phospho-AKT (Cell Signaling Technology, clone D9E). The following secondary antibodies were used: goat anti-rabbit Cy5 (Jackson Immunoresearch, 705-175-147), goat anti-mouse HRP (Thermo Scientific, PA1-28664). All primary antibodies were used at a 1:1000 dilution and secondary antibodies at a 1:10,000 dilution in 2% ECL Prime Blocking Reagent. The stripping protocol used was adapted from Yeung and Stanley, 2009. (Yeung and Stanley, 2009) Briefly, blots were washed 2 times with TBS-T, incubated twice for 5 minutes each in GnHCl stripping solution at room temperature (6 M GnHCl, 0.2% NP-40, 0.1 M β-mercaptoethanol, 20 mM Tris-HCl), washed 4 times (3 minutes each) in wash buffer at room temperature (0.14 M NaCl, 10 mM Tris-HCl, 0.05% NP-40), and then blocked in 2% ECL Primer Blocking Reagent in TBS-T. Stripped blots were re-probed following the protocol described.

### Mouse embryonic fibroblasts

Transformed mouse embryonic fibroblasts were cultured in (DMEM, 10% FBS, 1% L-glutamine, 1% penicillin/streptomycin, 1% HEPES, 1% sodium pyruvate) and used between passage 3-10. 1 x 10^4^ cells were seeded overnight onto poly-D-lysine coated glass coverslips in 24-well plates and then stimulated with 10 ng/mL recombinant mouse IL-1α (R&D), 10 ng/mL recombinant mouse TGF-β (R&D), or media alone for 48 hours. Coverslips were fixed in 4% paraformaldehyde for 10 minutes, permeabilized with 0.1% Triton X-100 for 15 minutes, blocked in 1% BSA for 30 minutes, and then stained overnight at 4°C with anti α-SMA antibody (Invitrogen, clone IA4). The next day, coverslips were stained for 1 hour at room temperature with phalloidin-eFluor660 (eBioscience) or donkey anti-mouse-AF594 (Jackson ImmunoResearch), and mounted onto slides with Vectashield Mounting Medium containing DAPI (Vector Laboratories). Coverslips were imaged on a Zeiss Imager M2 microscope (Carl Zeiss) with an AxioCam Mrm camera (Carl Zeiss) using a 20X objective (numerical aperture: 0.80) and ZenBlue software (Carl Zeiss). Cell area was determined by manually tracing cells in Fiji software. For serum starvation experiments, 2,500 MEFs/well were seeded into a 96 well plate overnight and then treated for 48 hours with cytokine in media containing either 10% or 1% serum. At the end of the 48-hour period, cell proliferation and viability were determined using CellTiter-Glo reagent (Promega).

### Flow Cytometry

Preparation of tissues for flow cytometric analysis was based on previously published protocols.(Mohar *et al*., 2015) Briefly, following euthanasia by CO_2_ and cardiac exsanguination at 9 weeks post-infection, mice were perfused through the heart with 10 mL perfusion buffer (HBSS, 5 mM HEPES, 0.5 mM EDTA), and liver, epigonadal visceral white adipose tissue, and quadriceps were excised. Following mechanical disruption of the tissue, tissue was collagenase-digested for 30 minutes at 37°C (shaking at 130 RPM) and filtered. After filtration, cells were centrifuged on a density gradient of 40% iodixanol, centrifuged 1038 x g for 25 minutes, no brake. Cells were washed and then blocked with anti-CD16/32 followed by staining with Zombie Aqua (BioLegend) and antibodies. The antibodies used for flow cytometry were: anti-F4/80-FITC (BioLegend, clone BM8), anti-Ly6C-PerCP/Cy5.5 (BioLegend, clone HK1.4), anti-CD19-PE/Cy7 (BioLegend, clone 6D5), anti-CD11c-APC (BioLegend, clone N418), anti-CD11b-AF700 (BioLegend, clone M1/70), anti-I-A/I-E-APC/Cy7 (BioLegend, clone M5/114.15.2), anti-CD45-PacBlue (BioLegend, clone 30-F11), anti-Ly6G-BV605 (BioLegend, clone 1A8), anti-CD3e-FITC (BioLegend, clone 145-2C11), anti-CD62L-PE-dazzle (BioLegend, MEL-14), anti-CD8a-PerCP/Cy5.5 (BioLegend, clone 53-6.7), anti-PD-1-PE/Cy7 (BioLegend, clone 29F.1A12), anti-CD4-APC (BioLegend, clone GK1.5), anti-CD44-BV605 (BioLegend, clone IM7). Stained cells were run on a BD Cytoflex flow cytometer and data was analyzed by FlowJo.

### qPCR on adipose tissue

Flash frozen tissues were homogenized in TRIzol reagent (Invitrogen) by bead-beating and RNA was extracted following manufacturer’s instructions. Following genomic DNA digestion and reverse transcription, RNA was run on a QuantStudio 6Flex (Applied Biosystems) using ABI Power SYBR Green PCR Master Mix (Applied Biosystems). Primers used: *ActB (Forward:* 5’-GGCTGTATTCCCCTCCATCG-3’, Reverse: 5’-CCAGTTGGTAACAATGCCATGT-3’), *Ucp-1 Forward: 5’-ACTGCCACACCTCCAGTCATT-3’ Reverse: 5’-CTTTGCCTCACTCAGGATTGG-* 3) *Prdm16 (Forward:* 5’-GAAGTCACAGGAGGACACGG-3’, *Reverse:* 5’-CTCGCTCCTCAACACACCTC-3’), *PGC1α (Forward: 5’-ACAGCTTTCTGGGTGGATTG-3’, Reverse:* 5’-TGAGGACCGCTAGCAAGTTT-3’), *Gdea (Forward:* 5’-TGCTCTTCTGTATCGCCCAGT-3’, *Reverse: 5’-GCCGTGTTAAGGAATCTGCTG-3), C/ebpβ (Forward:* 5’-TGACGCAACACACGTGTAACTG-3’, Reverse: 5’-AACAACCCCGCAGGAACAT-3’), *Pparγ-2 (Forward:* 5’-TCGCTGATGCACTGCCTATG-3’, Reverse: 5’-GAGAGGTCCACAGAGCTGATT-3’). Data were analyzed using the ΔCt method (relative to a housekeeping gene), and then normalized to the mean of the ΔCt of uninfected mice to get fold change.

### Body temperature measurements

Mice were anesthetized with isoflurane at 10 days post-infection, and subcutaneously injected with temperature micro-transponders (Bio Medic Data Systems, Seaford, DE). Temperature was monitored daily (at the same time of day) using a telemetric reader.

### Bomb calorimetry

Mice were individually housed for 24 hours at 10 weeks post-infection. Fecal pellets were collected every 2 hours during the daytime and flash-frozen. Pellets were lyophilized and then sent to University of Texas Southwestern Metabolic Phenotyping Core for bomb calorimetric analysis.

### CLAMS metabolic monitoring

At 5 weeks post-infection, mice were individually housed in Oxymax cages for 5 days (CLAMS, Columbus Instruments, Columbus, OH) at the University of Virginia. Mice were maintained at 25°C with 12 hour light/dark cycles and had free access to food and water in all conditions. The first 24 hours of data was considered and acclimation period and excluded from analysis. 16-point rolling averages across the remaining data were used to visualize data across the recorded time periods.

### Statistical analysis

Statistics were performed using GraphPad Prism 8. For comparison of two groups, two-tailed unpaired Student’s t-tests were performed with a confidence level of 95%. The Holm-Sidak test was used to correct for multiple comparisons. Statistical outliers were removed using the ROUT method (Q=1%), and this is indicated in the figure legend. Statistical significance threshold was set at P≤0.05. All data are presented as the mean ± SEM.

## Supplementary material

This manuscript has supplementary figures and an associated supplementary figure legend.

## Acknowledgements

We would like to thank Samantha Batista and Dr. Tajie Harris for sharing their initial observation that IL-1R^-/-^ regain weight after *T. gondii* infection, for assistance with the IL-1R immunofluorescence staining and critical feedback on the project. We thank Dr. Janet V. Cross for her critical evaluation of the manuscript. We thank Dr. Ping Hu for assistance preparing hydrogels and Riley T. Hannan for extensive discussions of fibroblast biology. We thank Dr. Ken Tung for histopathology advice. We thank Stefan Hargett for assistance with the CLAMS metabolic monitoring system and Marissa Gonzalez for assistance with flow cytometry preparation and analysis.

This work was supported by NIH K22 AI116727 (S.E.E.), T32 AI7496-23 (SM), NIH 1F32HL147405-01 (D.A)

The authors declare no competing financial interest.

## Author Contributions

S.J.M., J.A.H., and S.E.E designed the experiments in this study. Experiments were performed by S.J.M., J.A.H., E.A.L., C.M.S., K.A.B., I.S., and D.A., S. C-O. performed the blinded pathological assessment of histology. S.J.M. and S.E.E. prepared the manuscript.

