## Supplemental Figures for "Cachexia and fibrosis are costs of chronic IL-1R-mediated disease tolerance in *T. gondii* infection"

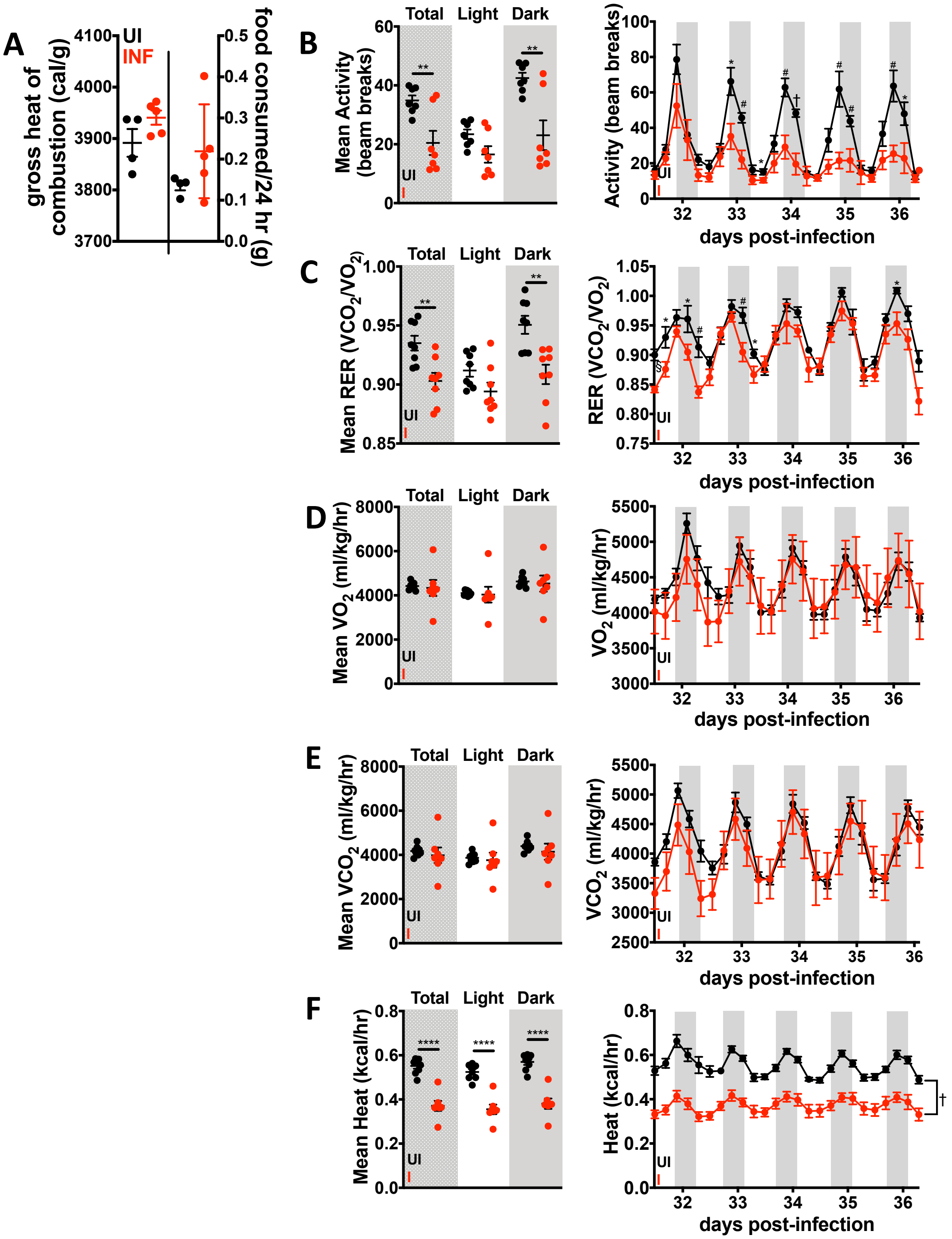


Supplemental Figure 1

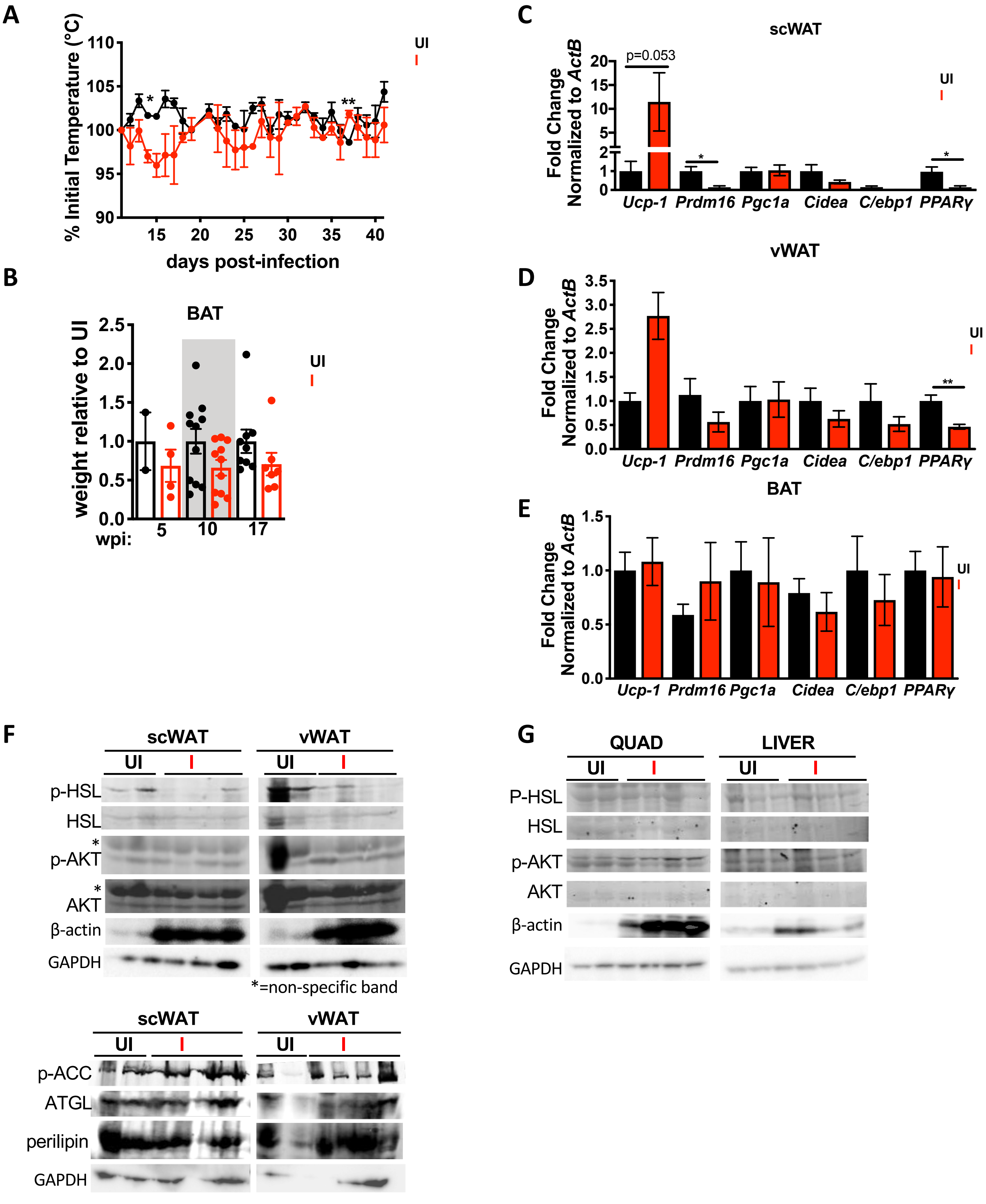


### Supplemental Figure 2

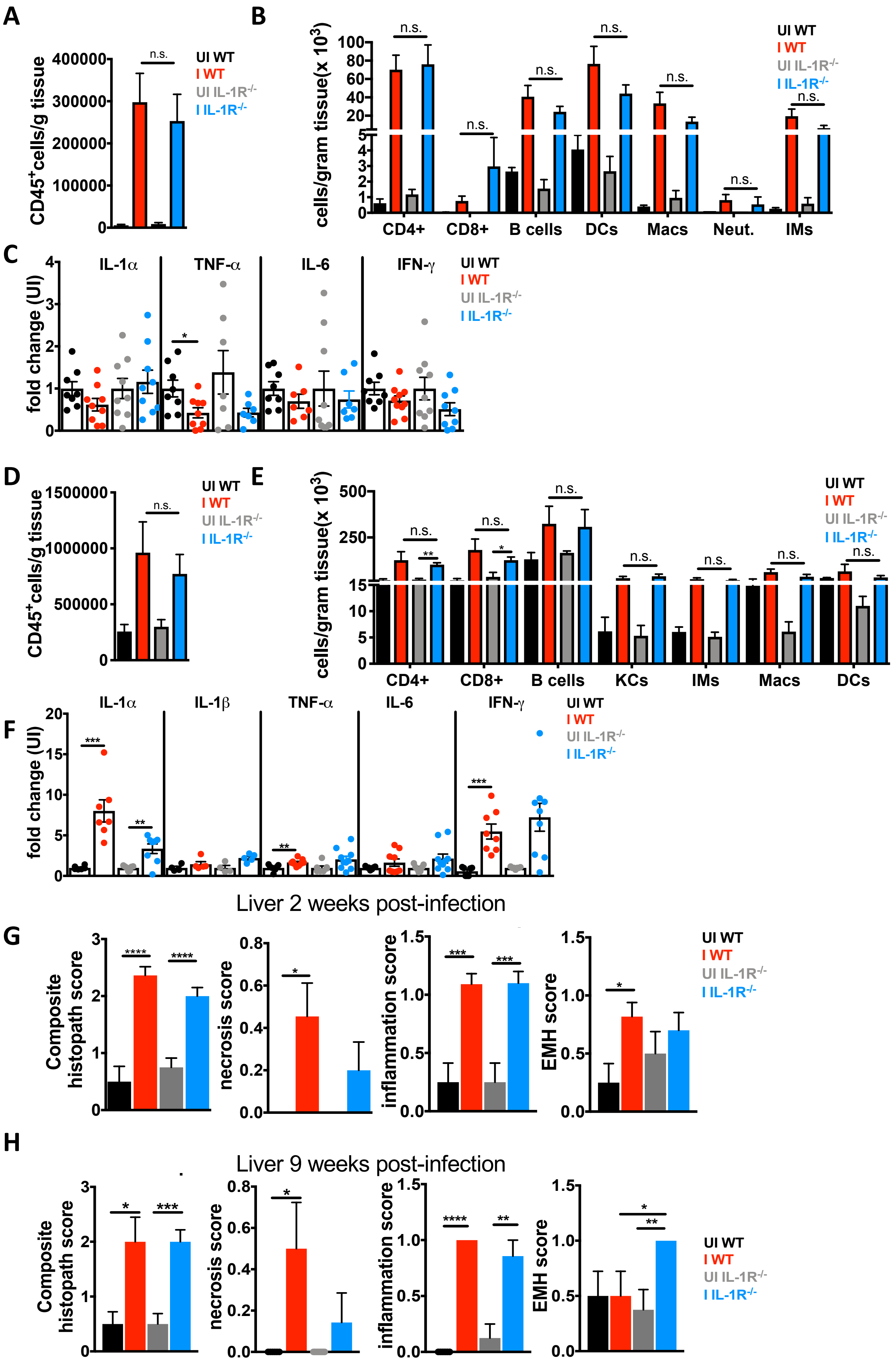


### Supplemental Figure 3

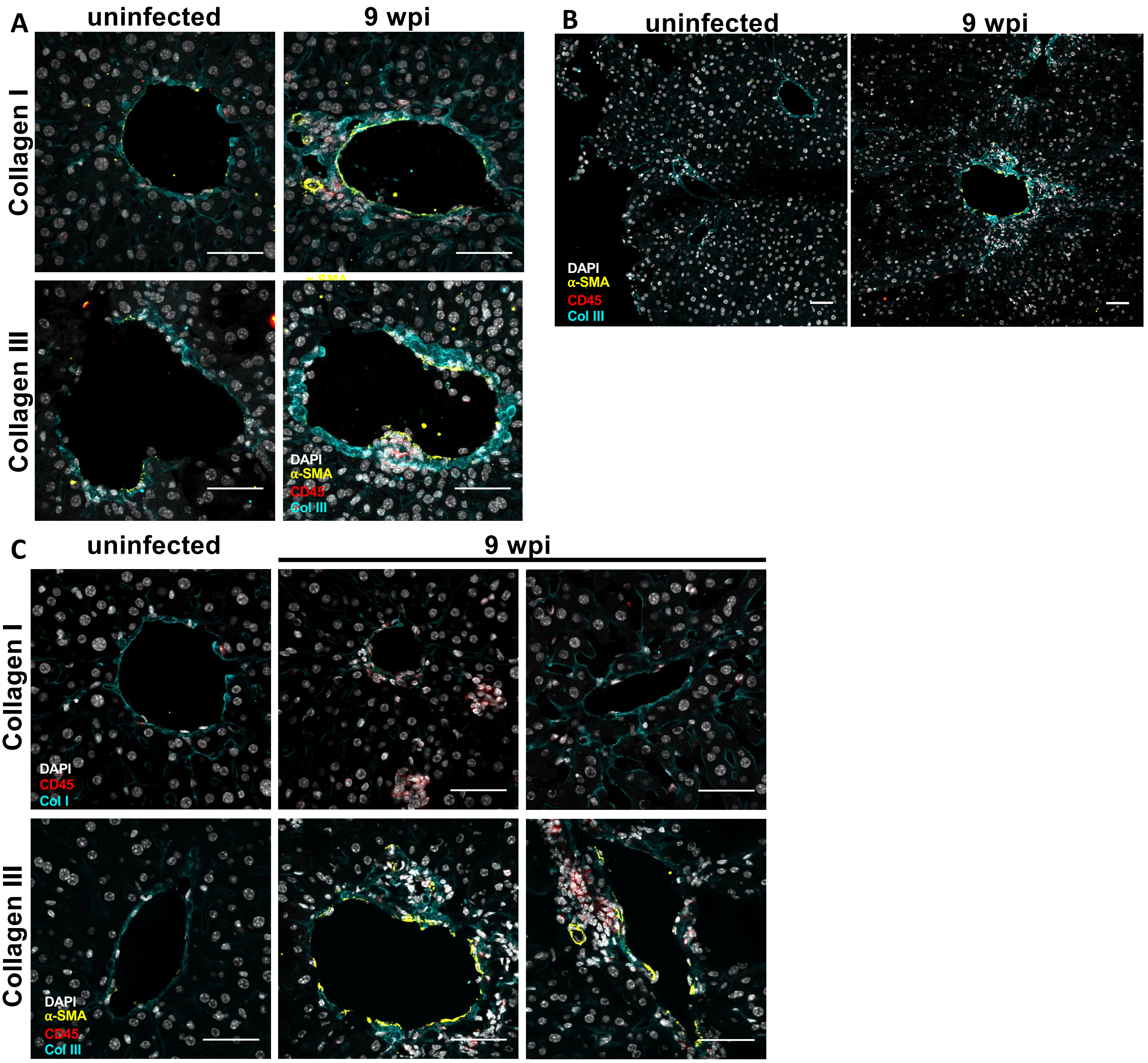


**Supplemental Figure 4**

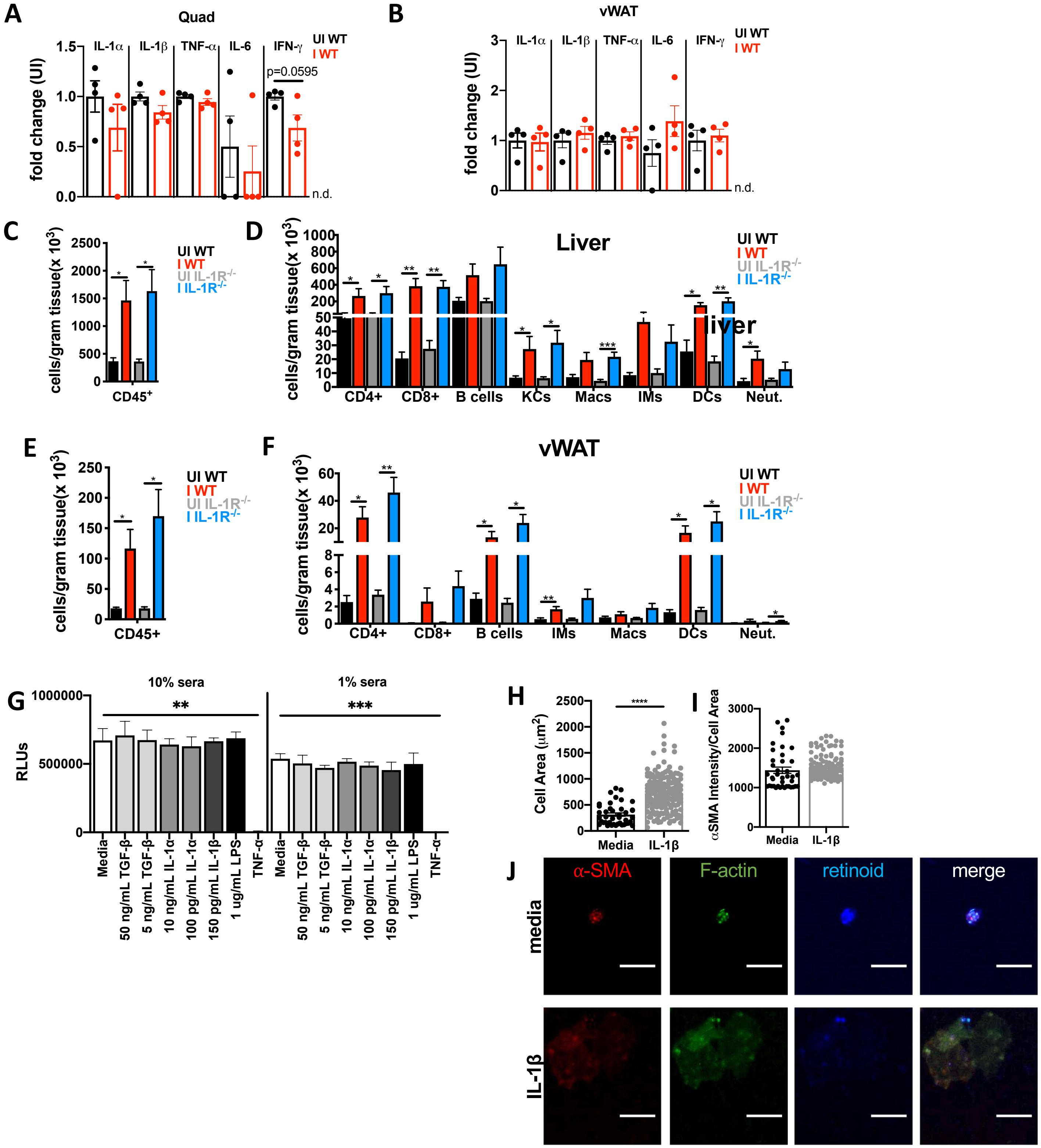


### Supplemental Figure 5

#### Supplementary Figure Legends

**Supplementary Figure 1: Cachectic mice have comparable food intake and nutrient absorption to uninfected mice. A,** 10-14 week old C57BL/6J mice (uninfected, UI and infected, I) were intraperitoneally infected with 10 Me49-GFP-luciferase *Toxoplasma* cysts and individually housed for 24 hours at 10 wpi. Caloric content of fecal pellets were determined by bomb calorimetry (left). Food intake over 24 hours was determined by weight (right). N= 5 mice per group. **B-F)** 10-14 week old C57BL/6J mice (uninfected, UI and infected, I) were intraperitoneally infected with 10 Me49-GFP-luciferase *Toxoplasma* cysts and housed in Oxymax CLAMS metabolic cages for the time points indicated. Data from the first 24 hours in the cage are excluded. Data on the left show mean of all the values from light (white bars) or dark cycles (shaded bars), or all time points combined (stippled bars). Data on the right show the 16 point rolling average of measurements with shaded bars representing the dark cycle over a 6 day span. **B,** Mean activity. **C,** Mean respiratory exchange ratio (RER). **D,** Mean $VO_2$ (normalized to lean body mass of the animal). **E,** Mean $VCO_2$ (normalized to lean body mass of the animal). **F,** Mean heat. N=7-8 animals per group, pooled between 2 independent experiments. Error bars are standard error of the mean*, $P < 0.05$; **, $P < 0.01$; ***, $P < 0.001$, by unpaired Student's T test with Holm-Sidak method to correct for multiple comparisons (mean data), and *, $P < 0.05$; $^{\#}$, $P < 0.01$; $^{\dagger}$, $P < 0.001$, $^{\S}$, $P < 0.0001$.

**Supplementary Figure 2: Non-shivering thermogenesis and lipolysis are not main drivers of *Toxoplasma*-induced cachexia.** 10-14 week old C57BL/6J mice (uninfected, UI and infected, I) were intraperitoneally infected with 10 Me49-GFP-luciferase *Toxoplasma* cysts. **A,** Mice were subcutaneously injected with telemetric temperature probes at 11 days post-infection, and temperature was monitored daily. **B,** Supraclavicular brown adipose tissue (BAT) weights relative to the mean weight of uninfected tissue at 5, 10, or 17 wpi. N=2-11 mice per group. **C-E,** qPCR for markers of fat browning and thermogenesis in inguinal subcutaneous adipose tissue (scWAT) **(C)** epigonadal visceral white adipose tissue (vWAT) **(D),** or supraclavicular brown adipose tissue (BAT) (**E**) at 17 wpi. Error bars are standard error of the mean. N= 7-9 mice per group. *$P < 0.05$; **$P < 0.01$; ***$P < 0.001$ by unpaired Student's T test with Holm-Sidak method to correct for multiple comparisons. **F-G,** Tissues were harvested at 5 wpi**.** Tissue lysates were made for subcutaneous white adipose tissue (scWAT) and epigonadal visceral white adipose tissues (vWAT) (**F**) or quadricep muscle (QUAD) and liver (**G**), and blotted for lipolysis machinery: phosphorylated and non-phosphorylated hormone sensitive lipase (p-HSL, HSL), phosphorylated and non-phosphorylated AKT, phosphorylated (p-ACC), perilipin. β-actin and GAPDH were loading controls. Each lane is lysate from an individual mouse. Representative of 2 independent experiments.

**Supplementary Figure 3. Liver and adipose tissue pathology is comparable between infected WT and IL-1R$^{-/-}$ mice at 2 and 9 wpi.** 12-14 week old C57Bl6 (WT, black UI, red I) or IL-1R$^{-/-}$ mice (gray UI, blue I) were intraperitoneally infected with 10 Me49 *Toxoplasma* cysts and vWAT **(A-C)** or liver **(D-H)** were harvested at 2 wpi or 9 wpi **(H). A,** total number of infiltrating CD45$^{+}$ immune cells per gram of vWAT at 2 weeks post-infection. **B,** CD4$^{+}$ T cells, CD8$^{+}$ T cells, B cells, dendritic cells (DCs), macrophages (Macs), neutrophils (Neut.), and inflammatory monocytes (IMs) in the vWAT at 2 wpi as determined by flow cytometry N=2-6 mice per group, pooled between two independent experiments. **C,** Cytokines in vWAT lysates 2 weeks post-infection were measured by ELISA. N=7-10, pooled from 2 independent experiments. **D,** total number of infiltrating CD45$^{+}$ immune cells per gram of liver at 2 weeks post-infection. **E,** CD4$^{+}$ T cells, CD8$^{+}$ T cells, B cells, dendritic cells (DCs), macrophages (Macs), neutrophils (Neut.), Kupffer cells (KCs), and inflammatory monocytes (IMs) in the liver at 2 weeks post-infection as determined by flow cytometry N=2-6 mice per group, pooled between 2 independent experiments. **F,** Cytokines in liver lysates 2 wpi were measured by ELISA. N=6-9, pooled from 2 independent experiments. Error bars are standard error of the mean. *P < 0.05; **P < 0.01; ***P < 0.001, n.s., not significant by unpaired Student's T test with Holm-Sidak method to correct for multiple comparisons. **G-H,** Liver was harvested at 2 wpi **(G)** or 9 wpi **(H)**, formalin fixed, and stained with H&E for histological analysis, including necrosis, inflammation, and extramedullary hematopoiesis (EMH). N=6-8 mice per group, pooled from 2 independent experiments.

**Supplementary Figure 3. Liver and adipose tissue pathology is comparable between infected WT and IL-1R$^{-/-}$ mice at 2 and 9 wpi.** 12-14 week old C57Bl6 (WT, black UI, red I) or IL-1R$^{-/-}$ mice (gray UI, blue I) were intraperitoneally infected with 10 Me49 *Toxoplasma* cysts and vWAT **(A-C)** or liver **(D-H)** were harvested at 2 wpi or 9 wpi **(H)**. **A,** total number of infiltrating CD45$^{+}$ immune cells per gram of vWAT at 2 weeks post-infection. **B,** CD4$^{+}$ T cells, CD8$^{+}$ T cells, B cells, dendritic cells (DCs), macrophages (Macs), neutrophils (Neut.), and inflammatory monocytes (IMs) in the vWAT at 2 wpi as determined by flow cytometry N=2-6 mice per group, pooled between two independent experiments. **C,** Cytokines in vWAT lysates 2 weeks post-infection were measured by ELISA. N=7-10, pooled from 2 independent experiments. **D,** total number of infiltrating CD45$^{+}$ immune cells per gram of liver at 2 weeks post-infection. **E,** CD4$^{+}$ T cells, CD8$^{+}$ T cells, B cells, dendritic cells (DCs), macrophages (Macs), neutrophils (Neut.), Kupffer cells (KCs), and inflammatory monocytes (IMs) in the liver at 2 weeks post-infection as determined by flow cytometry N=2-6 mice per group, pooled between 2 independent experiments. **F,** Cytokines in liver lysates 2 wpi were measured by ELISA. N=6-9, pooled from 2 independent experiments. Error bars are standard error of the mean. *P < 0.05; **P < 0.01; ***P < 0.001, n.s., not significant by unpaired Student's T test with Holm-Sidak method to correct for multiple comparisons. **G-H,** Liver was harvested at 2 wpi **(G)** or 9 wpi **(H)**, formalin fixed, and stained with H&E for histological analysis, including necrosis, inflammation, and extramedullary hematopoiesis (EMH). N=6-8 mice per group, pooled from 2 independent experiments.

**Supplementary Figure 5: Inflammatory infiltrate in the liver and vWAT is similar between infected WT and IL-1R$^{-/-}$ mice at 9 wpi. A-B,** 12-14 week old C57Bl6 (WT, black UI, red I) were intraperitoneally infected with 10 Me49 *Toxoplasma* cysts**.** Cytokines in quad lysates **(A)** or vWAT lysates **(B)** from mice 9 wpi were measured by ELISA. Data are presented as fold change relative to the mean of uninfected levels (N=4 mice per group). n.d. = not detectable. **(C, E)** Total number of infiltrating CD45$^{+}$ immune cells per gram of liver **(C)** or vWAT **(E)** at 9 weeks post-infection. **(D, F)** CD4$^{+}$ T cells, CD8$^{+}$ T cells, B cells, dendritic cells (DCs), macrophages (Macs), neutrophils (Neut.), Kupffer cells (KCs), and inflammatory monocytes (IMs) at 9 weeks post-infection were quantified in the liver (**D**) and vWAT (**F**)**.** N=6-7 mice per group, pooled between 2 independent experiments. Error bars are standard error of the mean. *P < 0.05; **P < 0.01; ***P < 0.001 by unpaired Student's T test with Holm-Sidak method to correct for multiple comparisons. **G,** MEF cells were plated on 96 well plates in 10% normal sera or 1% normal sera and incubated with media, IL-1α, IL-1β or TGFβ-1 for 48 hours. Cell number was determined relative to a standard curve using CellTiter-Glo reagent. Data pooled between 3 experiments. *P < 0.05; **P < 0.01; ***P < 0.001 by unpaired Student's T test. **H-J,** Primary hepatic stellate cells (HSCs) were isolated from uninfected mouse livers, and FACS sorted based on endogenous retinoid fluorescence. HSCs were seeded onto 4kPa hydrogels coated with 10ug/mL of fibronectin and cultured with 10ng/mL of IL-1b or media alone for 24hrs and then fixed and stained for F-actin and α-SMA and imaged by confocal microscopy. Cell area quantified in (**H**) and α-SMA intensity quantified as a proportion of cell area in (**I**). Error bars are standard error of the mean**.** ****P < 0.0001 by unpaired Student's T test. Statistical outliers were removed using the ROUT method (Q=1%). **J,** Representative images, scale bar represents 10 μm.
